# Correlative single molecule lattice light sheet imaging reveals the dynamic relationship between nucleosomes and the local chromatin environment

**DOI:** 10.1101/2023.11.09.566470

**Authors:** Timothy A. Daugird, Yu Shi, Katie L. Holland, Hosein Rostamian, Zhe Liu, Luke D. Lavis, Joseph Rodriguez, Brian D. Strahl, Wesley R. Legant

**Affiliations:** Department of Pharmacology, University of North Carolina at Chapel Hill, Chapel Hill, NC, USA; Joint Department of Biomedical Engineering, University of North Carolina at Chapel Hill, North Carolina State University, Chapel Hill, NC, USA; Janelia Research Campus, Howard Hughes Medical Institute, Ashburn, VA 20147, USA; Department of Biochemistry and Biophysics, University of North Carolina at Chapel Hill, Chapel Hill, NC, USA; National Institute of Environmental Health Sciences, Durham, North Carolina 27709, USA

**Author notes:** equal contribution (Timothy A. Daugird, Yu Shi).

**Keywords:** chromatin organization, epigenetics, microscopy, transcription, histone, nucleosome, phase separation, super-resolution

## Abstract

In the nucleus, biological processes are driven by proteins that diffuse through and bind to a meshwork of nucleic acid polymers. To better understand this interplay, we developed an imaging platform to simultaneously visualize single protein dynamics together with the local chromatin environment in live cells. Together with super-resolution imaging, new fluorescent probes, and biophysical modeling, we demonstrated that nucleosomes display differential diffusion and packing arrangements as chromatin density increases whereas the viscoelastic properties and accessibility of the interchromatin space remain constant. Perturbing nuclear functions impacted nucleosome diffusive properties in a manner that was dependent on local chromatin density and supportive of a model wherein transcription locally stabilizes nucleosomes while simultaneously allowing for the free exchange of nuclear proteins. Our results reveal that nuclear heterogeneity arises from both active and passive process and highlights the need to account for different organizational principals when modeling different chromatin environments.

## Introduction

The nucleus is a heterogeneous environment that is functionally and physically partitioned at multiple length scales. This partitioning occurs through spatial variations in the concentrations of nucleic acids and proteins within the nuclear space and allows the nucleus to perform diverse functions including DNA replication [1], transcription [2,3], RNA splicing [4], and ribosome biogenesis [5]. At the most fundamental scale, 147 DNA base pairs wrap around an octamer of histone proteins, collectively called a nucleosome [6]. Nucleosomes coalesce into heterogeneous groups of ∼4-15 clusters or ‘clutches’ at a scale of 10’s of nm [7,8]. These clutches further aggregate into irregular chromatin “nanodomains” containing 1000’s of base pairs that partition at the length scale of 100-200 nm and are separated by an interchromatin space that is enriched in RNA [9,10]. At the scale of the entire genome, individual chromosomes occupy distinct regions of the nucleus referred to as chromosome territories [11,12].

Computational and physical models provide insight to the thermodynamic principles by which chromatin organization is established (reviewed in [13]; however, a challenge remains to determine how variations in chromatin density within the nucleus relate to the functional specification of different nuclear processes. Single molecule imaging has shown that nucleosomes within nanodomain cores move coherently and are less mobile than those existing elsewhere in the nucleus [10,14]. Nucleosomes at the nuclear periphery, within dense chromocenters, or in differentiated cells (vs. pluripotent cells), all of which are enriched in heterochromatin, move less over time and are more radially confined than those in euchromatin [10,15]. Such findings suggest that dense heterochromatin regions may be more crowded and less physically accessible to nuclear proteins than sparse euchromatin regions [16]. Intriguingly, active processes appear to alter nucleosome motion in different ways. Transcription and DNA looping appear to stabilize nucleosomes [10,17,18] whereas DNA damage repair destabilizes them [17]. Together, these findings reveal a dynamic interplay between different nuclear processes, chromatin density, and the thermodynamically driven motion of nucleosomes. However, recent reports have also reported that nucleosome motion is independent of chromatin density, on average, when investigating replication inhibited cells with the same chromatin content, but double the nuclear volume compared to control cells [19] and other studies have demonstrated what even dense heterochromatin regions are equally accessible to an inert probe like green fluorescent protein [20].

Because of these discrepancies, we sought to better determine the relationship between chromatin density, nucleosome motion, the physical properties of the interchromatin space, and specific nuclear functions. Toward this goal, we developed an imaging platform that combines live-cell 3D single particle tracking (SPT) together with high-resolution volumetric imaging via lattice light sheet microscopy [21,22]. We used this platform to simultaneously visualize nucleosome motion in the context of local chromatin density variations in the nucleus. We combined these measurements together with fixed-cell 3D super-resolution imaging of nucleosome packing, diffusion measurements of inert fluorescent probes, targeted perturbations of specific nuclear functions, and biophysical modeling of chromatin organization.

Our findings demonstrate that nucleosome motion, spatial organization, and sensitivity to pharmacological and genetic perturbations all vary as a function of the local chromatin density in the nucleus. Intriguingly, the viscoelastic properties of the interchromatin space appear to be constant in both sparse and dense chromatin environments, suggesting that the observed differences in nucleosome behavior are more likely attributed to active processes, such as transcription, that locally stabilize nucleosomes in sparse euchromatic regions. Overall, our results provide a window into a heterogeneous and dynamic nuclear environment and provide an avenue to incorporate this heterogeneity into future models of chromatin function, spatial organization, and dynamics.

## Results

### Lattice light sheet microscopy enables simultaneous 3D tracking of individual nucleosomes together with high-resolution measurement of local chromatin density

We utilized a fibroblast-like Cos7 cell line that stably over-expresses histone H2b fused to a self-labeling HaloTag (Cos7-Halo-H2b) [23]. Previous reports indicate that exogenously expressed Halo-H2b is uniformly integrated into the mammalian genome [17] and co-labeling of our Cos-Halo-H2b cells with HaloTag-Ligand Janelia Fluor549 and Hoechst indicates that HaloTag-H2b integrates into sparse and dense chromatin with similar propensity (**Figure S1A**). Quantification via western blot indicated that Halo-H2b was expressed at roughly 4.4% compared to the endogenous protein (**Figure S1B**). To enable the simultaneous observation of individual nucleosomes and the surrounding chromatin microenvironment, we developed a two-color labeling and imaging protocol with lattice light sheet microscopy (**Figure1A, See methods**). We performed live-cell 3D single-molecule imaging with photoactivatable JaneliaFluor-646 to track nucleosomes with a lateral precision of 24 ± 9 nm and axial precision of 137 ± 59 nm (mean ± std, **Figure 1B, C, Figure S1C**) and simultaneous diffraction-limited volumetric imaging with JaneliaFluor 525 to resolve chromatin density with 334 x 837 nm resolution (full width half maximum) laterally and axially (**Figure 1C**). In this scenario, each voxel of the diffraction limited image could consist of approximately 90,000 base pairs of DNA and 625 nucleosomes. This assumes that nucleosomes are optimally packed within the volume (**Figure 1D**) and it is roughly comparable to the 500 nm estimate interaction radius of HiC [24].

**Figure 1:**
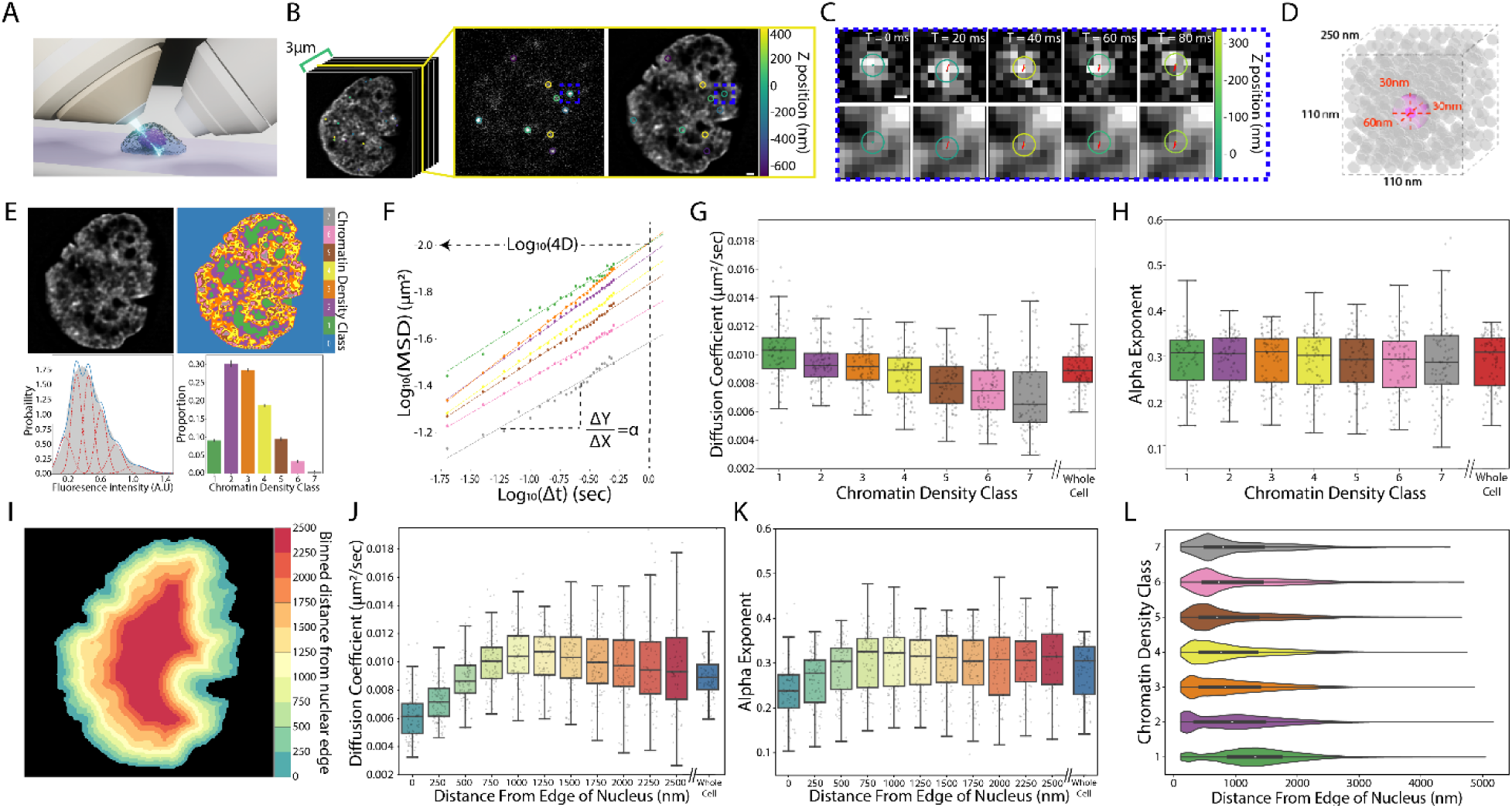
Nucleosome dynamics and their associated chromatin density. **(A)** Schematic of lattice light sheet microscopy imaging of chromatin dynamics. **(B)** A sample slice of single nucleosomes (middle) and their associated chromatin environment (right) from a 3D volume of cell nucleus (left). Circles indicate tracked nucleosomes, and the colors represent the z position relative to the objective focal plane. Scale bar =1000nm **(C)** The trajectory of the nucleosome in the blue box in (B). Top: Single nucleosome trajectory tracked across five consecutive 20 ms frames; Bottom: Overlay of nucleosome trajectory on the chromatin channel. **(D)** Schematic comparing LLSM voxel size (dashed cube), nucleosome localization precision (magenta sphere), and the potential nucleosome density (gray disks). **(E)** Chromatin density classification based on fluorescent intensity. Top left: deconvolved chromatin image; Top right: the corresponding chromatin density classification; Bottom left: histogram of chromatin intensity for top deconvolved image, the blue curves show the results for mixture of Gaussian fitting and red curves represent the individual components; Bottom right: histogram of the proportion of voxels within the top image that are assigned to different chromatin density classes. Error bars represent standard deviation of class proportions for the representative nucleus averaged across time points. **(F)** Representative mean square displacement (MSD) of nucleosomes in log-log scale for a single cell. The dashed lines show the linear fitting to a power law relationship (MSD = 4DΔt^α^) where α and D are the anomalous exponent and the apparent diffusion coefficient respectively. The colors indicate different chromatin density classes with the same convention as in (E). **(G)** Box plot of the extracted diffusion coefficient from (F) across different chromatin classes. The box indicates the inter-quartile range. The horizontal line in the middle indicates median, and the bars 1.5 x the upper and lower limits of inter-quartile range. Dots indicate fitted values averaged from a single cell. **(H)** Box plot of the extracted anomalous exponent α, the plot follows the same convention as (G). **(I)** Example distance to nuclear edge image generated for same cell as (E). **(J)** Box plot of the extracted diffusion coefficient as a function of distance from nuclear edge. The plot follows the same convention as (G). **(K)** Box plot of the extracted anomalous exponent as a function of distance from nuclear edge. The plot follows the same convention as (G). **(L)** Violin plot of the chromatin density classes and the distance from the nuclear edge of the voxels within each class. Data from G, H, J-K are from n = 88 cells across 8 independent biological replicates.

To isolate our SPT analysis specifically to DNA-bound H2b, we classified localizations as belonging to the same trajectory if they moved less than 400 nm over the course of sequential 20 ms frames. This approach allowed us to focus exclusively on the behavior of DNA-bound H2b and effectively filter out trajectories that may be due to free diffusing H2b or non-specifically bound dye molecules. To quantify chromatin density, we used an expectation maximization algorithm to segment the chromatin images into distinct classes based on intensity [25] (**Figure 1E**), hereafter referred to as chromatin density classes (CDCs). Partitioning chromatin into seven CDCs provided a robust fit to the intensity histograms while not overfitting the data according to the Bayesian information criterion and Akaike information criterion (**Figure S1D**). We found that simultaneous imaging of the underlying chromatin micro-environment was essential. Even at ∼2.5 sec/volume sampling, the underlying chromatin organization was quite dynamic with individual voxels transitioning through the entire range of CDC values within a time course (**Figure S1E, Movie S1**).

We note that these CDCs represent a statistical partitioning of a smooth intensity distribution rather than describing physically distinct chromatin regions. Nevertheless, the segmented chromatin images allowed us to quantify the relative amounts and spatial arrangement of dense vs. sparse chromatin in the nucleus and to partition nucleosome trajectories into similar chromatin densities across different cells.

### Single nucleosomes display differential motion based on local chromatin micro-environment

To investigate the relationship between chromatin density and nucleosome motion, we classified each nucleosome trajectory according to the underlying CDC and fit the cell ensemble mean square displacement (MSD) to a model of anomalous diffusion (**Figure 1F, Movie S2**). We found that, on average, nucleosomes in denser CDCs display a slower apparent diffusion coefficient (**Figure 1G**). This trend was consistent across eight biological replicates (**Figure S2A**), and showed significant differences in nucleosome motion across CDCs (Spearman coefficient = - 0.344, pvalue < 1E-5, pairwise t-tests in **Figure S2B**). Interestingly, we found that the negative correlation between chromatin density and nucleosome motion was strongest for the nucleosome diffusion coefficient and the radius of gyration (Spearman coefficient = −0.48, p < 1E-5) (**Figure 1G, Figure S2C**). Nucleosome displacement anisotropy displayed a smaller, though still significant correlation (spearman coefficient = −0.10, p = 0.006) (**Figure S2D**), while the anomalous diffusion exponent displayed no significant differences across CDCs (**Figure 1H**) (ANOVA F-statistic=0.31, p=0.93). This is further supported by analysis in which CDC’s were computationally scrambled in a manner that shifted their locations, but preserved their overall distribution and gross morphological features (**Figure S3A-C**). Under these conditions, we found no significant correlation between nucleosome dynamics and the spatially scrambled CDCs (ANOVA F-statistic = 0.21, p = 0.97) (**Figure S3D**). The observed differences in diffusion coefficients are notably larger than the combined precision of detection and tracking, as evidenced by the lack of correlation (ANOVA F-statistic = 0.24, p = 0.96) between the nucleosome diffusion coefficient and chromatin density class after paraformaldehyde fixation (**Figure S3D**).

We also observed a significant positive correlation between nucleosome diffusion coefficient and the distance from the nuclear edge in the outermost 1000 nm of the nucleus (t-test for the regression coefficient, p <0.001) (**Figure 1I, J**). Intriguingly, the alpha coefficient of nucleosome motion also shows a similar correlation within these regions, (t-test for the regression coefficient, p <0.001) (**Figure1K**), whereas it is unaffected by local chromatin density (**Figure1H**). To distinguish these effects on nucleosome motion from those due to chromatin density, we found that CDCs are distributed throughout the nucleus (**Figure1L**) with only a gradual trend of enrichment for high-density CDC’s near the nuclear periphery. Together, these findings demonstrate that nucleosomes in denser chromatin environments exhibit slower diffusion compared to those in sparser chromatin environments and indicates a link between chromatin density and nucleosome dynamics, with the diffusion coefficient and radius of gyration serving as key parameters.

We hypothesized that the observed differences in nucleosome motion across CDCs may arise from various factors, including variations in nucleosome spatial packing, differences in the local viscoelastic properties of the material surrounding chromatin, and/or differences in active processes that may be localized to a given CDC. These factors, either individually or in combination, could contribute to the distinct patterns of nucleosome dynamics observed in different CDCs. Subsequently, our investigation aimed to identify and characterize the biophysical properties and biological processes responsible for these observed differences in nucleosome motion.

### Nucleosomes in dense chromatin environments are more optimally space filling and randomly packed than nucleosomes in sparse chromatin environments

To measure the spatial arrangement of nucleosomes in different CDC’s we utilized 3D single molecule localization microscopy. Specifically, we labeled cells with a stochastically blinking HaloTag-JaneliaFluor630B probe [26], chemically fixed them with paraformaldehyde, and imaged them on a custom built highly inclined swept tile (HIST) microscope similar to that described in [27]. This arrangement allowed us to localize individual nucleosomes to 18 ± 8 nm laterally and 80 ± 28 nm axially (mean ± std, **Figure S4A**). To establish a comparable comparative framework to our live cell imaging, we convolved the 3D localization density histogram (**Figure 2B**) with an experimentally measured lattice light sheet point spread function, resulting in a simulated diffraction-limited image (**Figure 2C**). This simulated image then underwent the same processing pipeline and chromatin density classification as our live cell images (**Figure 2D, E**). Subsequently, each individual nucleosome localization was classified based on the underlying CDC (**Figure 2F, Movie S3**). Comparing the power spectrum of our simulated diffraction-limited images with that of our live cell images revealed quantitatively similar morphological features (**Figure S4B**). Furthermore, the distributions of the CDCs in our simulated images closely resembled those observed in our live cell imaging (**Figure S4C**). These findings indicate that our processing and classification pipeline successfully identifies and characterizes comparable chromatin density regions in both the super-resolution data and live cell imaging.

**Figure 2:**
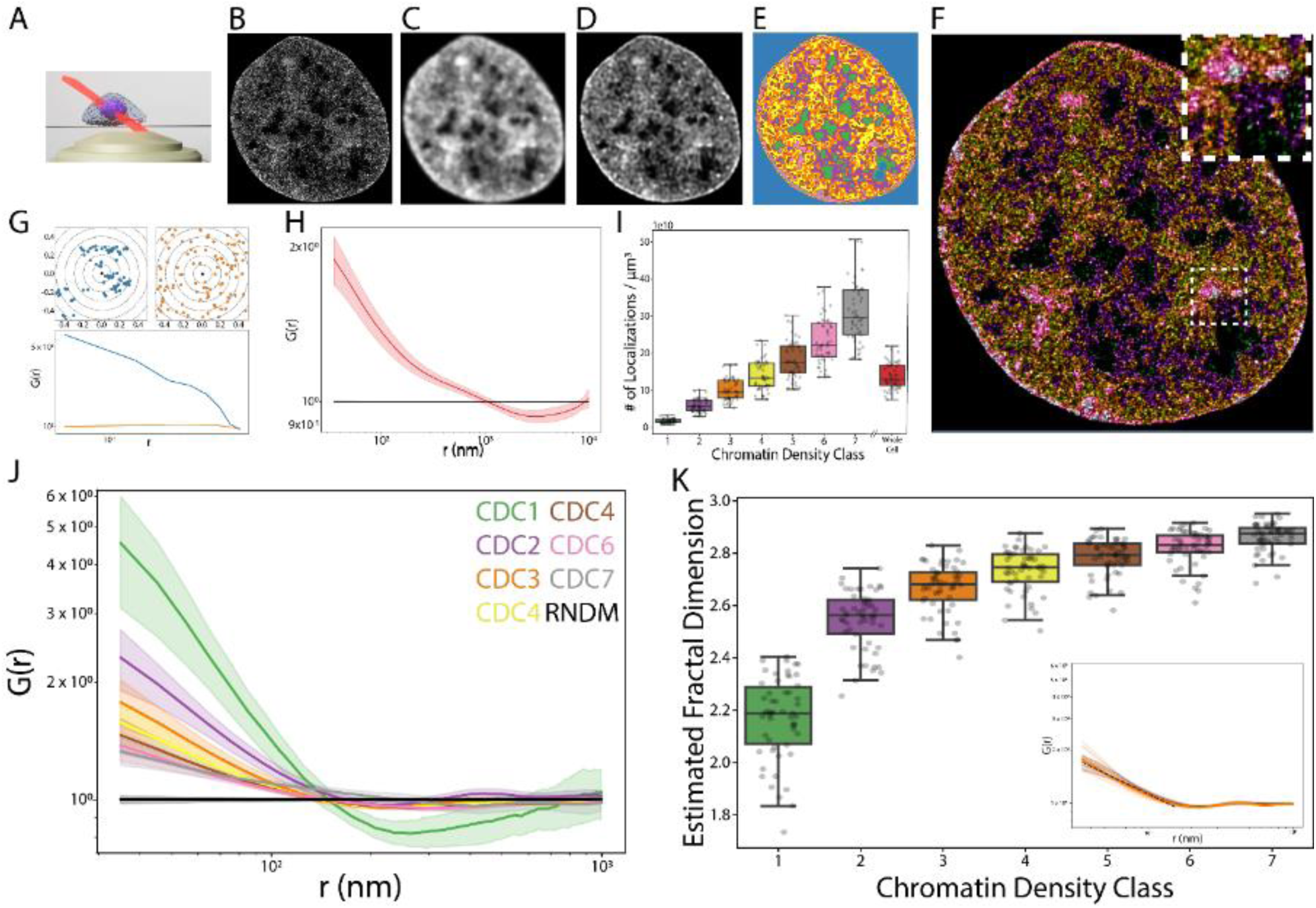
Measurements of nucleosome organization in different chromatin density classes. **(A)** Schematic of HIST microscopy imaging for single-molecule localization microscopy of nucleosome organization. **(B-E)** Representative images for the processing steps used to assign chromatin density classes to super-resolution nucleosome images. The super resolution reconstruction (B) is convolved with an experimental lattice light sheet microscopy point spread function to generate a simulated diffraction-limited image (C). This image is then deconvolved (D) and segmented into chromatin density classes (E). **(F)** Individual nucleosomes are assigned chromatin density classification according to their localization in (E). The top right dashed box in (F) is the zoom-in of the ROI – 2.3 x 2.3 μm. **(G)** Demonstration of the pair correlation function G(r) for a simulated clustered distribution (blue) and a simulated uniform distribution (orange). **(H)** Pair correlation function of the nucleosome organization for the entire cell (red) and a completely random distribution (black). The shade shows the standard deviation. **(I)** Box plot of localization density in different chromatin classes. The plot convention is the same as figure 1G. **(J)** Pair correlation function of the nucleosome organization for different chromatin classes. The black curve is the G(r) curve for a random distribution. **(K)** Box plot of the estimated fractal dimension for different chromatin classes. The plot convention is the same as figure 1F. Inset shows example power law fitting (black dashed line) to the G(r) curve for single chromatin density class in a single cell (Dark orange). Opaque lines represent G(r) curves for all other cells. Data from I-K are from n = 54 cells across 3 independent biological replicates.

To quantify nucleosome organization within each CDC, we computed the normalized 3D pair correlation function G(r), (**Figure 2G**) which quantifies the deviation from spatial randomness at a given length scale [28]. Unlike other methods for determining spatial clustering such as DBSCAN [29,30] or tesselation-based approaches [31,32], G(r) does not invoke implicit assumptions about discrete cluster sizes or cluster number. Furthermore, because G(r) is normalized by an equal number of randomly distributed points occupying the same space, it corrects for edge effects from masking and decouples the nucleosome spatial arrangement from variations in nucleosome density [28]. Consistent with previous reports [33], G(r) measurements of nucleosomes across the entire nucleus indicated linear scaling regimes, indicative of clustering along a continuum of spatial scales. The scaling can be characterized by two continuous power law like regimes: the first extends from the limit of our precision (∼30nm) to 186 nm ± 48 (mean ± std), and the second continues from there to 1088 ± 270 nm (**Figure 2H, Figure S4D**). This power law like behavior is consistent with a fractal-like organization of nucleosomes, with the power law exponent corresponding to a fractal dimension [34]. Fitting the linear regimes of our log-log plotted data yields estimated fractal dimensions of 2.72 ± 0.06 for the lower regime and 2.91 ± 0.02 for the upper regime (**Figure S4D**). These estimates are consistent with previous reports [33] and orthogonal methods for determining the fractal dimension of chromatin [20,35,36].

Interestingly, we found that, in addition to localization density (**Figure 2I**), the fractal dimension of nucleosome packing was also positively correlated with chromatin density (Spearman correlation coefficient = 0.78, **Figure 2J, K**). The lowest density CDC 1 had a fractal dimension of 2.14 ± 0.25 whereas the highest density CDC 7 had a fractal dimension of 2.85 ± 0.09. The largest jump was observed between CDCs 1 and 2 (t-statistic =-23.89, p < 1E-5) with a progressive increase in fractal dimension between CDCs 2 through 7 (Spearman coefficient =0.67, p < 1E-5). When we repeated this analysis in scrambled CDCs, we found no clear differences in localization density (ANOVA F-statistic = 0.94, p = 0.46 or G(r) curves (**Figure S4E,F**). These deviations from random organization were also clearly seen in representative reconstructions of the nucleosome localizations in different CDCs (**Figure S5**).

Overall, these results demonstrate that the spatial arrangement of nucleosome packing varies together with the local chromatin density in the nucleus. Nucleosomes in low density CDCs display a lower fractal dimension, are more clustered, and less randomly distributed than nucleosomes in high density CDCs which are more randomly distributed and more optimally space filling.

### The interchromatin space displays similar viscoelastic properties regardless of chromatin density

We next sought to investigate the relationship between chromatin density and the viscoelastic properties of the interchromatin space. To this end, we extended our two color labeling and imaging protocol to track the motion of a non-interacting HaloTag fused to a nuclear localization signal (HaloTag-NLS) as it diffused within the nucleus (**Movie S4**). For each CDC, we calculated the cell-ensemble MSD for free-diffusing HaloTag-NLS and extracted the anomalous exponent and apparent diffusion coefficient through fitting a power law model (**Figure 3A-C**). In contrast to our nucleosome measurements, we found no apparent dependence in either the apparent diffusion coefficient or the alpha parameters on chromatin density classes. This indicates that, for a non-interacting free diffusing particle of ∼2.6 nm size [37], different chromatin density classes display similar viscoelastic properties. We also found that the anomalous exponent is about 0.8, which is close to what one would expect for diffusion in purely viscous liquid (anomalous exponent of one).

**Figure 3:**
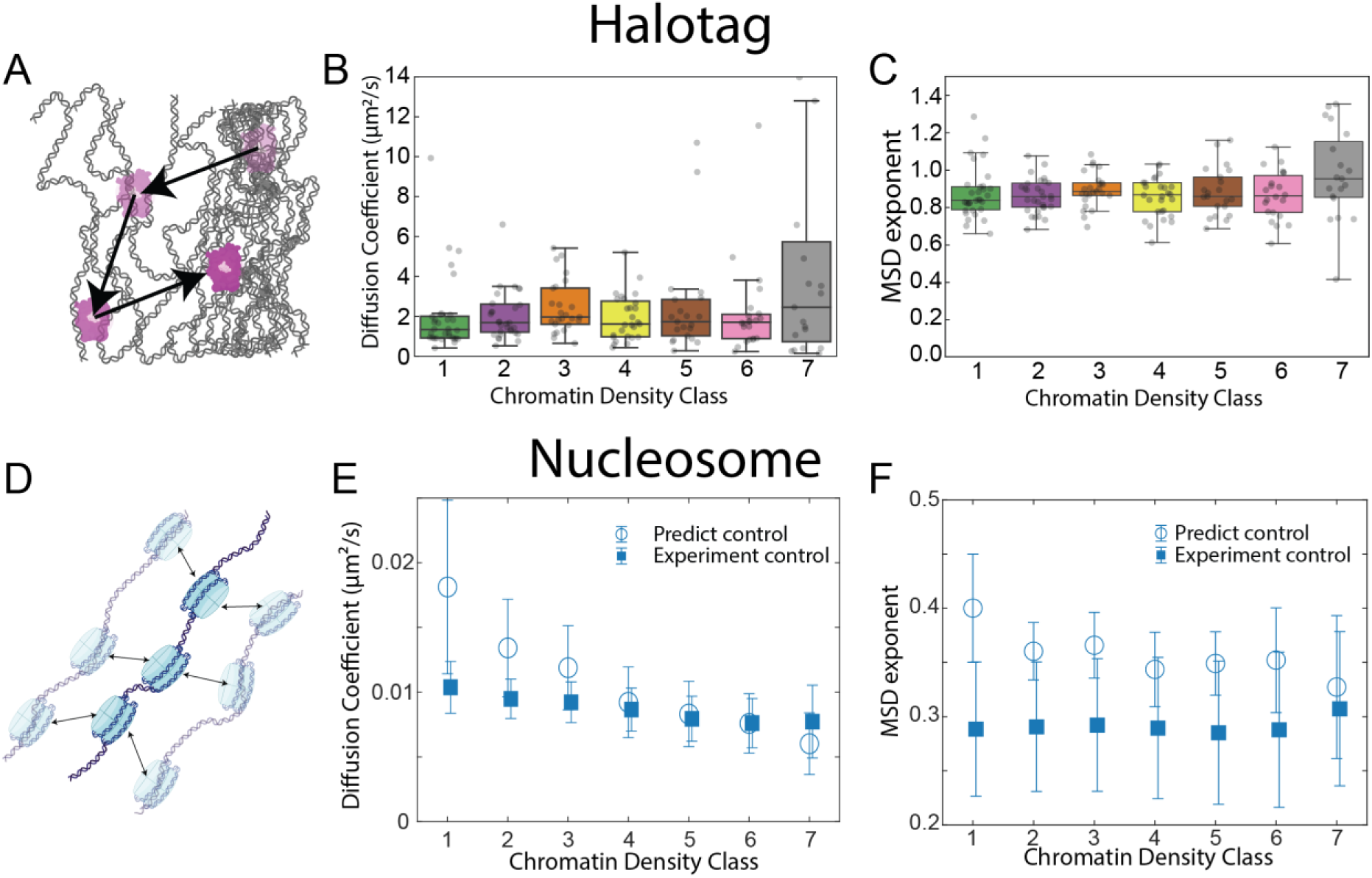
Measurement of nucleus viscoelasticity using free HaloTag-NLS. **(A)** Schematic of free HaloTag-NLS (magenta) diffusing through the chromatin fiber network (black lines). **(B)** Box plot of the extracted diffusion coefficient for HaloTag-NLS across different chromatin density classes. The figure convention is the same as Figure 1G. **(C)** Box plot of the extracted anomalous exponent, α, for HaloTag-NLS across different chromatin density classes. The figure convention is the same as Figure 1G. **(D)** A schematic of nucleosome motion diffusion. **(E)** The comparison between the model-predicted and the experimentally measured (Figure1G) nucleosome diffusion coefficients across different chromatin density classes. Open circles indicate model predicted results and closed squares indicate experimentally measured results. The error bars represent the standard deviation of the experimental data. For the simulations, error bars are computed by propagating the error from the experimentally measured terms to the model output. **(F)** The comparison between model-predicted and experimentally measured (Figure1G) nucleosome MSD exponent across different chromatin classes. The figure convention is the same as (E). Data from B and C are from n = 37 cells across 4 independent biological replicates.

### Biophysical models can integrate nucleosome motion, density, and packing properties

To integrate these results together with the nucleosome spatial organization and diffusive behavior observed above, we sought to extend existing biophysical models of nucleosome motion count for our analysis of chromatin density classes. Previous studies have modeled chromatin as a Rouse polymer comprised of nucleosomes that passively diffuse within a viscoelastic medium [38,39]. We specifically chose the model described in [39] because it incorporates the fractal organization of nucleosomes, the viscoelastic properties of the medium and has an analytical form for the predicted nucleosome MSD that can be compared to our experimental observations. As predicted in the model, the nucleosome MSD can be expressed as *MSD*(*t*) = *D_*app*_t^*β*^*, with 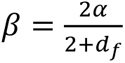 and 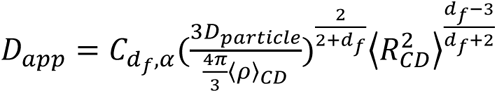. In these equations, 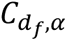 is a dimensionless parameter determined from the chromatin fractal organization and the viscoelastic properties of the interchromatin space (see Methods). *α* and *D*_*particle*_ are the anomalous exponent and apparent diffusion coefficient of a freely diffusing particle within the interchromatin space, and *d*_*f*_, ⟨*ρ*⟩_*CD*_, and 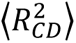 are the fractal dimension, the nucleosome density, and the chromatin domain size respectively.

To compare this model to our observations, we estimated *α* and *D*_*particle*_ from our free-diffusing HaloTag-NLS measurements and combined these with estimates of *d*_*f*_, ⟨*ρ*⟩_*CD*_, and [inine] from our nucleosome measurements within each CDC (see Methods for details). We then compared the model-predicted nucleosome apparent diffusion coefficient and anomalous exponent within each CDC to our experimentally measured values (**Figure3D-F, Figure S6A-E**). Overall, the model-predicted values for the nucleosome diffusion coefficient and the anomalous exponent were in a similar range to our experimental measurements. The model also recapitulated a decrease in nucleosome diffusion coefficient with increased chromatin density (**Figure3E**) with only a modest dependence of the anomalous exponent on chromatin density (**Figure3F**). Surprisingly though, the predicted sensitivity of the diffusion coefficient to chromatin density was much stronger than what we observed experimentally. At dense CDCs (CDC>=4), the predicted nucleosome diffusion coefficient largely overlapped with our experimental measurements (predicted 0.0068±0.0023 μm^2^/s vs. experimental 0.0077±0.0024 μm^2^/s for CDCs 6-7), but the model substantially overestimated the nucleosome diffusion in sparse chromatin classes (predicted 0.0158±0.0052 μm^2^/s vs. experimental 0.0099±0.0018 μm^2^/s for CDCs 1-2).

Together, these results suggest that a fractal Rouse polymer model that incorporates the viscoelastic properties of the internuclear space can accurately predict nucleosome dynamics in dense chromatin regions, but that there appear to be additional factors that lead to slower than predicted nucleosome motion in sparse regions. This next led us to investigate what might be responsible for stabilizing nucleosomes within the low density CDCs.

### Pharmacological perturbations alter nucleosome dynamics and suggest a partitioning of nuclear functions between chromatin density classes

Previous reports have found that proteins related to active processes, such as transcription and DNA looping, are predominantly found in areas of sparse and intermediate chromatin density [9]. For these reasons, we hypothesized that the discrepancy between model and experimental data could be due to active biological processes locally stabilizing nucleosome motion in sparse chromatin regions. To test this hypothesis, we employed a panel of pharmacological perturbations to inhibit various steps of active transcription (**Figure 4A**). Consistent with previous reports [10,17,18], our findings revealed that treatments including α-amanitin (aAM), 5,6-Dichloro-1-β-D-ribofuranosylbenzimidazole (DRB), and Pladienolide B (PladB) resulted in accelerated nucleosome dynamics throughout the entire nucleus (**Figure S7**). Notably, the largest difference in nucleosome motion between the transcription inhibitors and control groups was observed in CDC1-2 (**Figure 4B**). These differences decrease in magnitude through the intermediate classes, finally dropping below the level of significance in dense CDCs. The effects of these perturbations on nucleosome dynamics were primarily evident in the apparent diffusion coefficient, with no clear effects on the anomalous diffusion exponent (**Figure S7B**). Consistent with these results, in HCT116 cells in which the major sub-unit of RNA polymerase II, RPB, has been engineered to be conditionally knocked down in the presence of the drug 5PH-I-AA [40] (**Figure S7C**), we see comparable, albeit less class specific, effects on nucleosome dynamics (**Figure S7D,E**).

**Figure 4:**
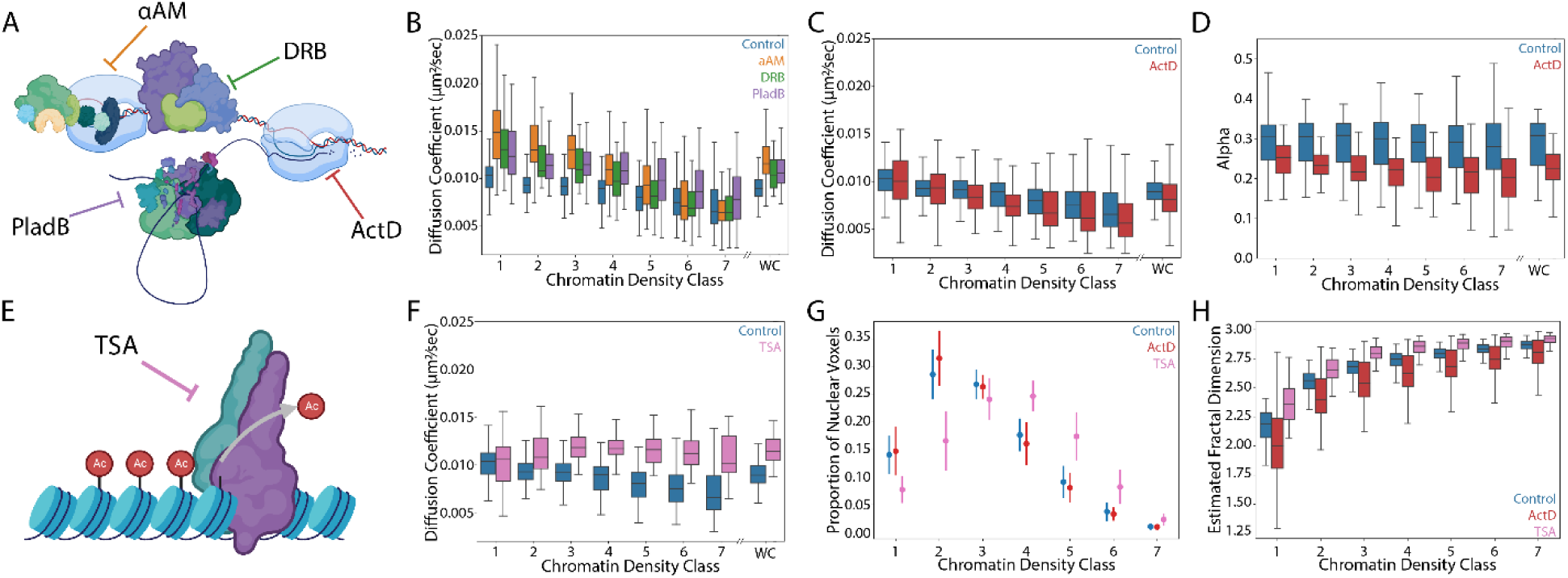
Measurements of the relationship between nuclear functions and nucleosome properties. **(A)** Schematic of different perturbations targeting gene transcription. **(B)** Box plot of diffusion coefficient in different chromatin density classes under control (blue), α-amanitin (orange), DRB (green) and PladB (purple). WC stands for whole cell. **(C, D)** Box plot of diffusion coefficient and anomalous exponent in different chromatin density classes under control (blue), actinomycin D (red). **(E)** Schematic of Trichostatin A (TSA) perturbation on histone deacetylase activity. **(F)** Box plot of diffusion coefficient in different chromatin density classes under control (blue) and TSA (pink). **(G)** Point plot displaying the distribution of nuclear voxels in different chromatin classes under control (blue), ActD (red) and TSA (pink) conditions. The central dot indicates the mean proportion of voxels in given class for all cells. The vertical line indicates the standard deviation. **(H)** Box plot of estimated fractal dimension from super-resolution images of different chromatin classes under control (blue), ActD (red) and TSA (pink) conditions. Data from B, C, D, F, and G are from n = 88 cells across 8 replicates (control), n = 52 cells across 3 replicates (α-amanitin), n = 46 cells across 3 replicates (DRB), n = 49 cells across 3 replicates (PladB), n = 41 cells across 3 replicates (ActD) and n = 60 cells across 3 replicates (TSA). Data from H are from n = 54 cells (control), n = 54 cells (ActD), and n = 60 cells (TSA) across 3 replicates. All replicates above were independent biological replicates.

Contrary to the effects observed with other pharmacological perturbations of transcription, but consistent with previous findings [17], we found that actinomycin D (ActD) treatment resulted in reduced nucleosome dynamics throughout the nucleus (**Figure S7A**), and across all CDCs (**Figure 4C**). We also observed consistently more anomalous diffusion, as indicated by a decreased anomalous diffusion exponent in all CDCs upon ActD treatment (**Figure 4D**). Importantly, ActD operates through a distinct mechanism compared to the other tested perturbations. Unlike the other transcriptional perturbations, ActD does not act directly on the protein machinery but rather intercalates DNA, thereby hindering RNA polymerase from transcribing along the DNA template. Based on this mechanism, we propose that ActD induces a global “stiffening” effect on the DNA polymer. This hypothesis is supported by previous in vitro studies demonstrating ActD’s ability to alter DNA persistence length and contour length, ultimately resulting in increased stiffness [41].

To determine whether perturbations of other nuclear functions, beyond just transcription, might affect nucleosome motion in a chromatin-density dependent manner, we next investigated the effects of the broad-spectrum histone deacetylase inhibitor Trichostatin A (TSA). Previous studies have shown that TSA relaxes chromatin spatial organization [42,43] and increases average nucleosome dynamics across the nucleus [10]. Here, we show that TSA treatment increased nucleosome motion to the greatest extent in dense chromatin classes and abolished the relationship between nucleosome diffusion coefficient and CDC (ANOVA stat = 1.45, p=0.19; **Figure 4F**). TSA treatment did not induce changes anomalous diffusion exponent (**Figure S7F**).

These data establish a compelling link between previous reports examining the relationship between chromatin density and specific markers of transcription in fixed cells and nucleosome dynamics in live cells and support the hypothesis that active transcription locally stabilizes nucleosomes in low-density CDCs [17]. To confirm this, we sought a more direct means of visualizing active transcriptional processes in live cells by monitoring the transcriptional bursting of endogenously regulated genes using the MS2 system in cells that were co-labeled with the DNA intercalating SiR-Hoechst dye (**Movie S5**). We employed a previously established MCF7 cell line that stably expresses GFP-tagged MS2 (GFP-MCP) coat protein and contains 24x MS2 stem loops integrated into the 3’ end of the endogenous loci of the estrogen response TFF1 gene [44]. When TFF1 is transcribed, GFP-MCP concentrates to the MS2 RNA stem loops, appearing as a bright green spot. Consistent with our results suggesting that transcription primarily occurs in sparser chromatin regions, we found that the initiation of TFF1 transcription bursts primarily occurred in CDC1-2, with a smaller proportion in CDC3-4 **(Figure S8)**.

### Pharmacological perturbations have differential effects on nucleosome dynamics and nucleosome organization

Unlike the distinct trends that link nucleosome dynamics with chromatin density classes upon transcription inhibition, inhibiting either the formation of the transcription pre-initiation complex using aAM or transcription elongation using DRB had no significant impact on nucleosome organization. This was evident at both the mesoscale, where no noticeable shift in the proportions of chromatin density classes was observed in our live-cell imaging (**Figure S7G**), and at the nanoscale, indicated by the consistent distribution of fractal dimensions across all classes similar to control cells (**Figure S7H**). Interestingly, we found that the spliceosome inhibitor PladB induced a small, though significant change in the proportional representation of CDCs in the nucleus in our live cell imaging (**Figure S7G**), as well as the apparent fractal dimension (**Figure S7H**). Previous studies indicate that ActD leads to increased chromatin compaction[45–47], whereas treatment with TSA is associated with chromatin relaxation [42,48]. While we did not observe significant changes in the proportional representation of CDCs following ActD treatment (**Figure 4G**), we did find a more clustered nucleosome organization at the nanoscale. This was evidenced by a decrease in estimated fractal dimensions across all CDCs (**Figure 4H**). Conversely, TSA appeared to have a ‘normalizing’ effect on chromatin organization at both the mesoscale and the nanoscale. At the mesoscale, TSA treatment resulted in a distribution of CDCs that was more Gaussian (**Figure 4G**). At the nanoscale, TSA led to a more random organization of nucleosomes, as indicated by an increase in fractal dimensions across all CDCs (**Figure 4H**).

### Pharmacological perturbations do not alter viscoelastic properties of the interchromatin space with respect to small and inert proteins

It is possible that the observed effects of pharmacological perturbations on nucleosome dynamics might be explained by indirect changes to the viscoelastic properties of the interchromatin space (e.g. through decreased amounts RNA produced) rather changes to chromatin itself. To test this, we measured the diffusive motion of non-interacting HaloTag-NLS under each of the pharmacological perturbations described above. Interestingly, although the perturbations did increase or decrease HaloTag-NLS motion across the nucleus as a whole when compared to control cells, none of the perturbations had a CDC-specific effect on HaloTag-NLS motion (**Figure S9**). Finally, we tested how these perturbations to different nuclear functions affected the agreement between our experimentally observed results and the biophysical model described above (**Figure 5, Figure S10**). Interestingly, transcription inhibitors aAM and DRB yielded better agreement between experimental results and model predictions for the nucleosome diffusion coefficients and the MSD exponents, especially in low-density CDCs. This suggests that gene transcription plays a role in actively stabilizing nucleosome motion within lower density CDCs in a manner that is distinct from the purely thermal forces considered in the model. We also found that TSA treatment led to the model performing worse in dense CDCs. This is likely because the model does account for electrostatic nucleosome-nucleosome interactions that are altered under TSA-induced hyper acetylation. These results highlight the need to account for both active (non-thermal) forces as well as passive (thermal) forces when modeling chromatin dynamics.

**Figure 5:**
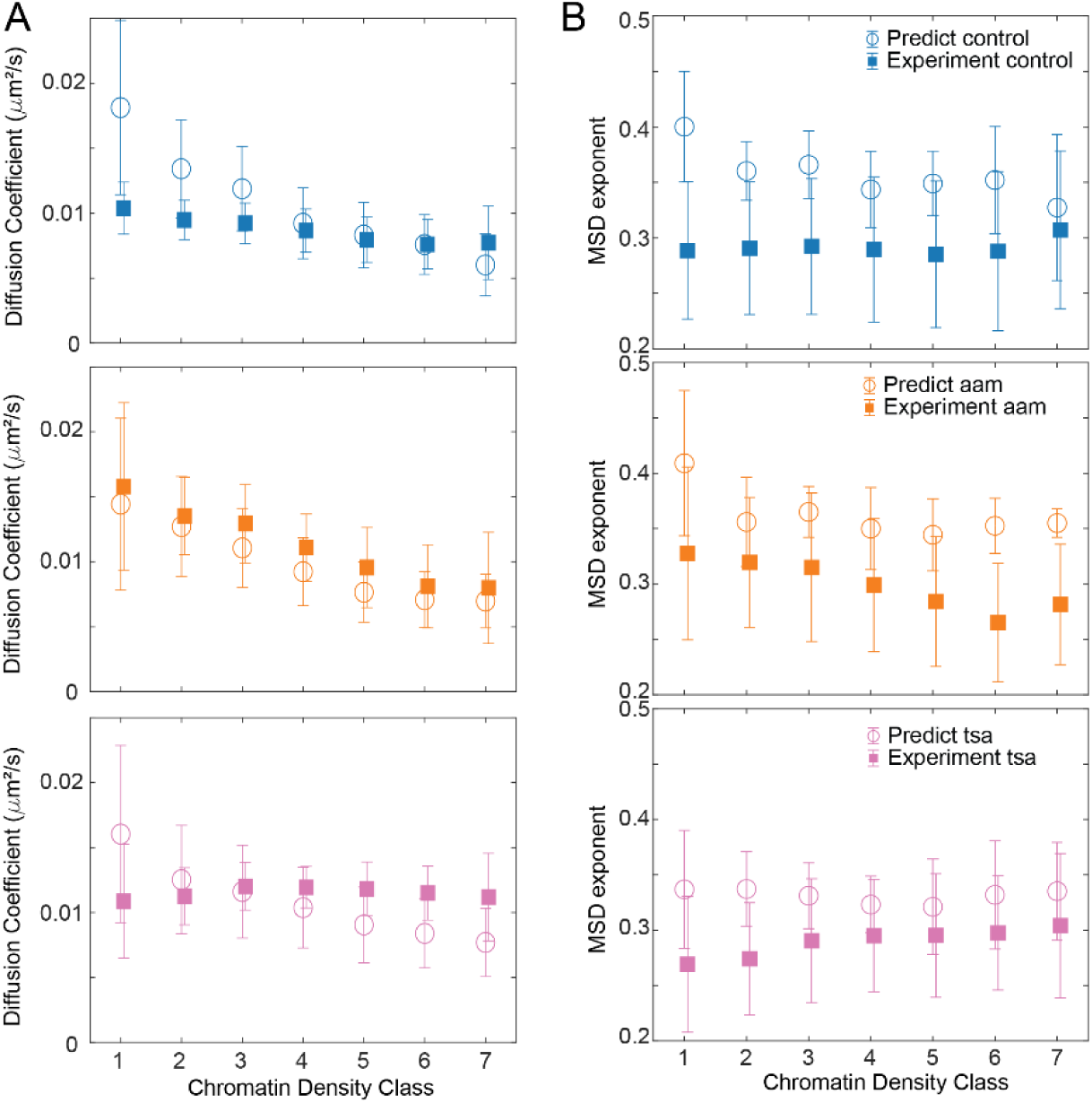
Comparisons between model predicted and experimentally measured nucleosome dynamics under different pharmacological perturbations. **(A)** Comparison of diffusion coefficients between model (open circles) and experiments (closed squares) under control (blue), α-amanitin treatment (orange), and TSA treatment (pink). Error bars indicate standard deviations. **(B)** Comparison of MSD exponents between model and experiments under control (blue), α-amanitin treatment (orange), and TSA treatment (pink). The plot convention is the same as (A).

## Discussion

A variety of biochemical reactions occur simultaneously within the nucleus of each cell. How these reactions are coordinated in absence of partitioning biological membranes is an active and ongoing area of investigation [49] that has been aided by technological developments in live cell [50], single molecule [10,15,17] and super resolution [8,9,42] imaging as well as by theoretical modeling efforts in polymer sciences [13,39,51,52]. Here, we add to this field by developing a new imaging approach to track individual nucleosomes and free diffusing proteins with nm precision while simultaneously reporting the local density and viscoelastic properties of the surrounding chromatin microenvironment at a resolution of a few hundred nanometers. With this methodology, we revealed that nucleosome motion is negatively correlated with local chromatin density. This finding agrees with and expands upon prior studies that explored nucloeosome motion by binarizing the nucleus into heterochromatin vs euchromatin regions [15], but is distinct from reports of decreased nucleosome motion at the nuclear periphery [10]. While decreased nucleosome motion has previously been suggested to be due to increased heterochromatin at the nuclear periphery [10], it is important to note that the laminal region of the nucleus possesses additional distinct characteristics beyond chromatin density (reviewed in [53]). Specifically, the laminal DNA is physically tethered to the inner nuclear membrane [54,55], generally lacks active gene regulatory processes [56,57], and is enriched for distinct histone post translational modifications [56,58]. Here, we separately differentiate between the local chromatin density around a nucleosome and the nucleosome’s distance from the nuclear periphery. Intriguingly, we observe that the alpha coefficient of nucleosome motion is positively correlated with the distance from the nuclear periphery, but is unaffected by local chromatin density. These findings suggest that there are likely additional effects, beyond changes to chromatin density, which impact nucleosome motion near the nuclear periphery. Compared with previous work that focused on either the dynamics [10,46] or the spatial organization [8,9,33,59] our approach provides an integrated view to investigate both properties in 3D simultaneously.

We complemented these measurements with fixed-cell, 3D super-resolution microscopy using a new generation of spontaneously blinking Janelia-fluor probes and found that nucleosome packing is more structured and clustered in sparse chromatin regions and becomes more optimally space filling and randomly organized in dense chromatin environments. Nucleosome packing displays a fractal-like organization between a length scale of ∼40-400 nm, the lower-bound of which is imposed by our averaged 3D localization error. This length scale falls within the range of previously reported chromatin nanodomains [9,10,59], but is larger than the reported size of nucleosome clutches in which small groups of nucleosomes organize into discrete clusters [8,15]. This apparent discrepancy can be explained by our observation that the fractal dimension of nucleosome packing varies as a function of chromatin density, ranging from 2.1 in sparse CDCs to 2.8 in dense CDCs, a range which is comparable to that reported previously with variety of different methods [20,33,35,59]. A lower fractal dimension suggests that nucleosomes in sparser CDCs are more frequently organized into small concentrated regions (such as clutches) separated by nucleosome free regions. In dense CDCs, as the inter-nucleosome spacing begins to approach the ∼8 nm length scale of the nucleosome itself [60], the packing becomes more optimally space filling which results in a more random organization and a higher fractal dimension. These trends were also visually apparent from our super-resolution images **(Figure S5)**.

Interestingly, the interchromatin space displayed predominantly viscous properties regardless of chromatin density, at least when probed by an inert HaloTag molecule. This indicates that from the perspective an inert nuclear protein of comparable size to a transcription factor, the nucleus is predominantly liquid-like and that this is independent of local chromatin density. This is consistent with earlier results measuring the viscoelastic properties of cell nucleus using quantum dots or fluorescent dextran [20,61] and indicates that even dense heterochromatin regions are still highly accessible. We want to emphasize that our delineations between chromatin dense and chromatin sparse CDCs only account for nucleosome density and do not report information about the concentrations of other nuclear proteins or nucleic acids. In fact, both chromatin-dense and chromatin-sparse CDCs can be highly crowded with other biomolecules. The fact that the dynamics of freely-diffusing HaloTag show little dependence on local chromatin density suggest that the effective pore size for a diffusing nuclear protein of similar size to HaloTag is relatively homogeneous across the entire cell nucleus. The simple explanation for this is that regions with lower chromatin density are more crowded with other biomolecules and visa-versa (**Figure 6**, [45])

**Figure 6.**
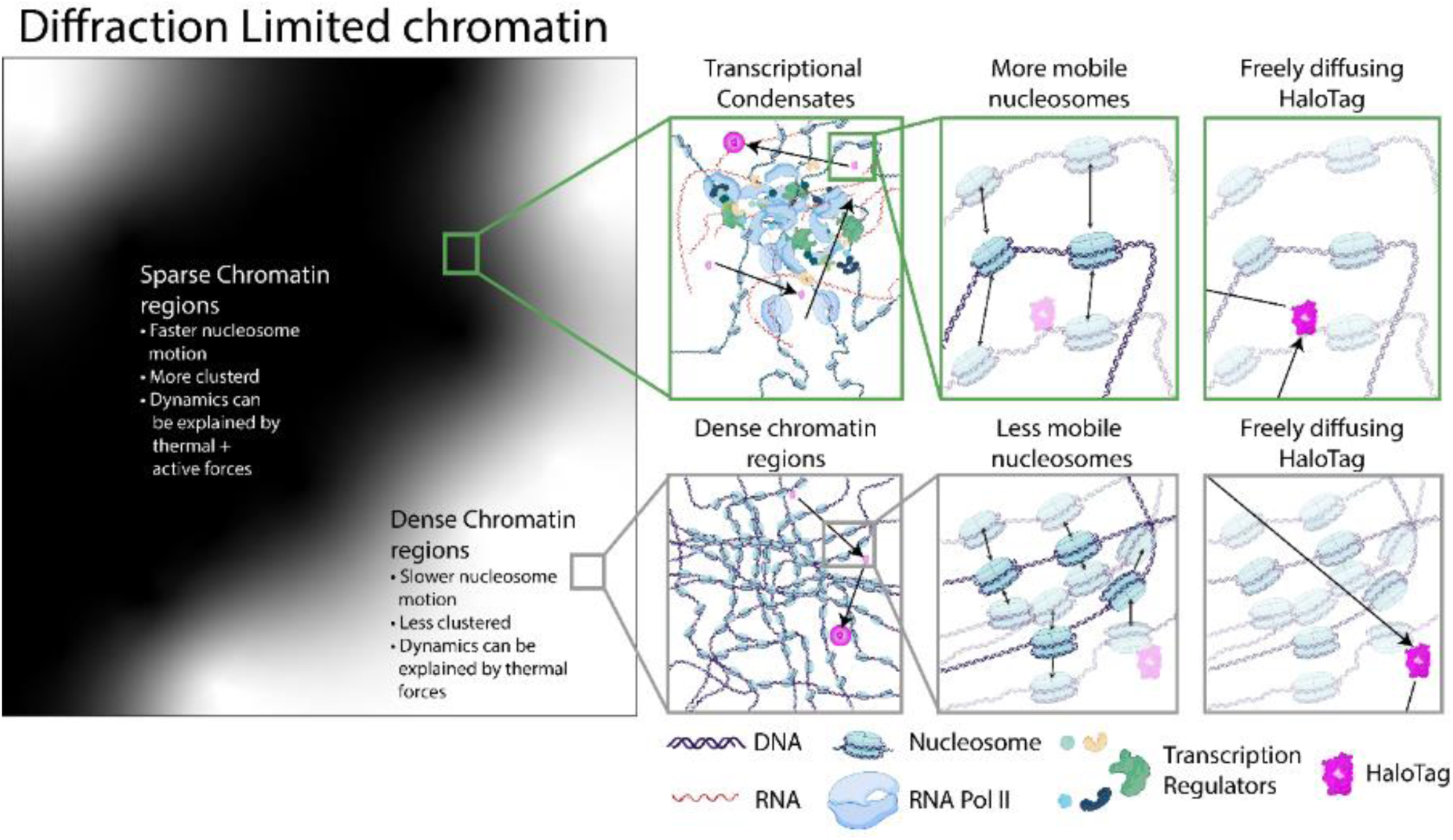
Proposed model for chromatin density and organization. Dark regions in the left panel represent chromatin sparse regions (green box). Nucleosomes in these regions form small clusters that are separated by nucleosome free regions and are more mobile than nucleosomes in denser chromatin regions. Their motion can be explained by a combination of passive diffusion and an active stabilizing component due to gene transcription. Bright regions represent chromatin dense regions (gray box). Nucleosomes in these regions are more randomly organized and display less motion than those in chromatin sparse regions. Their motion can predominantly be explained by passive diffusion. In contrast, the diffusion of inert particles (magenta) is not affected by the local chromatin density (right column) or perturbations to transcription.

There have been several elegant attempts to connect polymer models [15,39,51,62] to experimentally measured nucleosome dynamics and spatial organization [15,35]. Our results complement these efforts by highlighting the heterogeneity in dynamics, organization, and biological function of chromatin and the challenges when treating the cell nucleus as a single material. For instance, a universal fractal globule model with a single fractal dimension may not fully describe the chromatin spatial organization [13]. Rather, we show that the apparent fractal dimension depends on the local chromatin density. The heterogeneity in chromatin dynamics and organization suggests similarities between the cell nucleus and colloidal glasses [63,64] and offers a possible explanation for how chromatin can actively undergo reorganization while maintaining its structural integrity. Moreover, biological processes such as gene transcription play a larger role in regulating nucleosome motion in chromatin sparse regions than chromatin dense regions. It will be important to include this heterogeneity, together with the role of active process that may locally affect nucleosome dynamics in specific nuclear regions in future physical models of the cell nucleus.

Previous efforts to connect nucleosome dynamics to biological processes have produced conflicting results. Tracking single nucleosomes in RPE cells or single gene loci via an inserted MS2/TetR system revealed decreased chromatin motion during active gene transcription [17,65]. In contrast, tracking of several loci using dCAS targeted promoters and enhancers showed that chromatin motion was increased in settings wherein the genes were actively transcribing [66]. At the whole-nucleus level, our pharmacological perturbations agree with the model wherein transcription stabilizes nucleosome motion. Moreover, our finding that this stabilization predominantly occurs in low chromatin density regions, favors the model that transcriptional hubs with a higher local concentration of transcription regulatory machinery act to stabilize the local chromatin environment [17]. Our finding that this stabilization affects nucleosomes, but not the diffusion of free HaloTag molecules suggests several possible mechanisms. Chromatin stabilization could be accomplished through specific chemical interactions e.g. between bromodomain containing proteins within the transcription hub to acetylated histone tails of the nucleosomes. Alternatively, it could arise from the generation of a local meshwork of RNA, DNA, and transcription machinery that has a pore size that is larger than single transcription factors, yet smaller than single nucleosomes. In this manner, nuclear region could be stabilized in a transcriptionally competent state while allowing for free exchange of smaller and dynamic transcription factors and supporting machinery.

In summary, our work reveals the relationship between nucleosome dynamics, organization, nuclear viscoelasticity, and biological processes. Our findings support the need to account for physical and molecular heterogeneity in future biophysical models and will inform future studies of for how chromatin spatial organization and dynamics regulate cell development [67], disease [68,69], and stem cell reprogramming [8,15,70]. We envision that extensions of our imaging approach to simultaneously visualize single molecules within the context of their local microenvironment will be useful in future studies to explore the relationship between chromatin density, nuclear function, and the diffusion of a diverse spectrum of nuclear proteins.

## Acknowledgements

We thank John Crocker for helpful feedback and advice and Rachel Cherney, Mauro Calabrese, and Max Hockenberry for assistance with biochemical and imaging experiments. We also thank Victoria Augoustides for assistance in figure preparation. This work was funded in part by grants from the National Institutes of Health (1DP2GM136653) awarded to W.R.L.. W.R.L. acknowledges additional support from the Searle Scholars program, the Beckman Young Investigator Program, and the Packard Fellowship for Science and Engineering.

## Author Contributions

W.R.L. conceived the project together with T.A.D. Y.S. and Z.L. L.D.L and K.L.H. generated the fluorescent Janelia-Fluor probes used in this study. H.R. and B.D.S. assisted with biochemical characterization of cell lines and protein expression quantification. T.A.D., Y.S., and W.R.L. performed the imaging experiments, analyzed the data, and wrote the manuscript with feedback from all authors. W.R.L. supervised and directed the project.

## Methods

### Sample preparation

#### Cell culture

Cos7-Halo-H2b and Cos7-Halo-NLS cells were maintained in DMEM (Thermo Fisher Scientific, 11965118), supplemented with 10% fetal bovine serum (Avantor 1300-500H) and 100 units/mL penicillin-streptomycin (Thermo Fisher Scientific, 15140122). HCT116 cells were maintained in McCoy’s 5A (Thermo Fisher Scientific 16600082) supplemented with 10% fetal bovine serum (Avantor 1300-500H) and 100U/mL penicillin-streptomycin (Thermo Fisher Scientific, 15140122). MCF7-TFF1-MS2 cells were maintained in MEM (Thermo Fisher Scientific 11090081) supplemented with 10% fetal bovine serum (Avantor 1300-500H), 2mM Glutamine (Thermo Fisher Scentific, 25030081) and 100 units/mL penicillin-streptomycin (Thermo Fisher Scientific, 15140122). Cells were routinely checked for mycoplasma contamination through both imaging and isothermal PCR methods.

#### Generation of Halo-H2b cell lines

Cos7-Halo-H2b cells were a generous gift of Luke Lavis. HCT116-mClover-mAID-RPB [40] cells were a generous gift of Dr. Masato Kanemaki at the National Institute of Genetics, Japan. Cos7-Halo-NLS and HCT116-mClover-mAID-RPB+HaloTag-H2b cell lines were generated using piggyBac transposon-based methods. The cDNA for Halo-NLS and HaloTag-H2b_BsdR genes was synthesized by GeneScript, then ligated into piggyBac plasmids that conferred resistance to either gentymycin or blasticidin [71,72]. Around 10^5^ Cells were plated in 6 well plates and allowed to recover for 24 hours. Cells were transfected with 1500ng of HaloTag-NLS or HaloTag-H2b + 1000ng of piggyBac Transposase (System Biosciences, PB210PA-1) using Lipofectamine2000 (Thermo Fisher Scientific, 11668027) and PLUS reagent (Thermo Fisher Scientific, 11514015) according to manufacturer’s protocol. Two days post transfection, cells were harvested and plated into a 25 cm^2^ flask in selection media. For Cos7-Halo-NLS cells, the selection media was DMEM (Thermo Fisher Scientific, 11965118), supplemented with 10% fetal bovine serum (Avantor 1300-500H) and 800µg/mL of Geneticin (Thermo Fisher Scientific, 10131027). For HCT116-mClover-mAID-RPB+Halo-H2b cells, selection media was HCT116 growth media supplemented with 6µg/mL of blasticidin (GoldBio, B-800-25).

#### Co-localization of HaloTag-H2b and Hoechst sample preparation

Cos7-HaloTag-H2b cells were plated on a 25 mm, #1.5 coverslips at a density of approximately 10^4^ cell/cm^2^ and allowed 24 hours to recover from resuspension. Cells were incubated with 100 nM of Halo Tag Ligand Janelia Fluor 647 (HTL-JF647) for 1hour, washed with pre-warmed PBS (Thermo Fisher Scientific, 10010023) then incubated in fresh growth media for 20 minutes. 4% Paraformaldehyde solution was prepared through combining 10mL of 16% PFA (Electron Microscopy Sciences 15710), 4 mL of 10x PBS (Corning 45001-130), and 26 mL of diH2O. Cells were washed briefly with PBS and fixed in room temperature 4% PFA for 12 minutes. Fixed cells were washed once with PBS for 5 seconds, followed by three five minute washes in PBS (10x dilution of (Corning 45001-130). All fixed samples were washed according to this washing scheme unless otherwise noted. Cells were incubated with 10µg/mL of Hoechst (Thermo Fisher Scientific, H3570) for 20 minutes then washed according to previously described PBS washing scheme. Cells were mounted on 20 x 75 x 1.0 mm slide using vectashield (Vector Laboratories, H-1700-2) and sealed with nail polish 1 hour later.

#### Labeling of HaloTag-H2b cells for live cell imaging

One day prior to imaging, cells were plated on 25 mm, #1.5 coverslips at a density of approximately 10^4^ cells/ cm^2^. On the day of imaging, the cells were incubated in cell culture media containing 10 nM of the Halo tag ligand Photo-activatable Janelia Fluorophore 647 (HTL-PA-JF647) for 1 hour. The cells were then washed with pre-warmed PBS (Thermo Fisher Scientific, 10010023) and incubated for an additional 20 minutes in Cos7 or HCT116 growth media containing 100 nM of either Halo tag ligand Janelia Fluorophore 525 for Cos7-HaloTag-H2b cells (HTL-JF525) or 100 nM of Halo tag ligand Janelia Fluorophore 549 (HTL-JF549) for HCT116-mClover-mAID-RPB+HaloTag-H2b cells. After the incubation, the cells were washed with pre-warmed PBS (Thermo Fisher Scientific, 10010023) and incubated in cell culture media containing no dye for 10 minutes.

#### Labeling of cells for super resolution imaging

Two days prior to imaging, cells were plated on 24 well #1.5 glass bottomed plates (Cellvis, P24-1.5H-N) at approximately 10^5^ cells/cm^2^. One day prior to imaging, cells were incubated in growth media supplemented with 500nM Halo tag ligand Janelia Fluor 630b for one hour. Cells were then washed with pre-warmed PBS (Thermo Fisher Scientific, 10010023) and cells were incubated in fresh growth media for 20 min. A 250 ng/mL solution of wheat germ agglutinin conjugated fluorescent nanodiamonds (Adámas, NDNV1000-WGA custom order) in PBS (10x dilution of Corning 45001-130) was sonicated for at least 1 hour. Cells were washed 1x room temperature PBS (10x dilution of Corning 45001-130), and fixed in freshly prepared, room temperature 4% PFA for 12 minutes. Post fixation, cells were washed once with PBS (10x dilution of Corning 45001-130) for 5 seconds, followed by three five minute washes in PBS (10x dilution of Corning 45001-130). Cells were then incubated in the sonicated NDNV100nm-WGA solution solution for 1 hour to allow for fiducial nanodiamonds to bind to the cell. Cells were washed once with PBS (10x dilution of Corning 45001-130) for 5 seconds, followed by three five minute washes in PBS (10x dilution of Corning 45001-130). Cells were imaged either immediately after washing or stored at 4° C for no more than 48 hours.

#### Labeling of Cos7-HaloTag-NLS cells for live cell imaging

One day prior to imaging, cells were plated on 25 mm, #1.5 coverslips at a density of approximately 10^4^ cells/ cm^2^. On the day of imaging, the cells were incubated in cell culture media containing 10 nM of the 647 (HTL-PA-JF647) and a 500x dilution manufacturer recommend stock concentration of SPY505 (Cytoskeleton, CY-SC101) for 1 hour. After the incubation, the cells were washed with pre-warmed PBS (Thermo Fisher Scientific, 10010023), and incubated in Cos7 growth media containing no dye for 10 minutes.

#### Pharmacological perturbations

The following concentrations of drugs were used for pharmacological perturbations, α-Amanitin: 100 ug/mL (Millipore-Sigma A2263-1MG); 5,6-dichloro-1-beta-D-ribofuranosylbenzimidazole Sigma-Aldrich D1916): 100 µM; Actinomycin D (Millipore-Sigma A9415-2MG): 0.5 µg/ml; Pladienolide B (Tocris, 6070) 30 ng/mL; Trichostatin A (Sigma-Aldrich T8552-5MG) 300 nM; 5PH-I-AA (MedChemExpress HY-134653): 1 µM. For pharmacological inhibition of transcription and RNA splicing with the drugs above, cells were incubated with the appropriate drug for 30 minutes prior to dye labeling. During dye labeling, the labeling and wash solution was also supplemented with the appropriate concentration of the drug. The total time of drug incubation for each of these conditions prior to imaging was 2 hours. For pharmacological inhibition of histone deacetylases with Trichostatin A, cells were plated two days prior to imaging. Cells were incubated with Trichostatin A for 16 hours prior to imaging. The dye and wash solutions were also supplemented with Trichostatin A. The total time of TSA incubation prior to imaging was 18 hours. All live cell imaging was performed in Fluorobrite DMEM (Thermo Fisher Scientific A1896701), supplemented with 10% fetal bovine serum (Avantor 1300-500H), 2mM Glutamine (Thermo Fisher Scentific, 25030081), 100 units/mL penicillin-streptomycin (Thermo Fisher Scientific, 15140122), and containing the drug concentrations listed above.

#### MS2 sample prep

TFF1-MS2 cells were described previously[44]. Transcription of the estrogen response gene was induced as previously reported. Cells were plated on 25 mm, #2 coverslips at least three days prior to imaging. At least two days prior to imaging, growth media was replaced with hormone depletion media, consisting of pheonol free MEM (Thermo Fisher Scientific, 5120003), supplemented with 10% Charcoal stripped FBS (Millipore-Sigma F6765), 2mM Glutamine (Thermo Fisher Scentific, 25030081) and 100 units/mL penicillin-streptomycin (Thermo Fisher Scientific, 15140122). Media was replaced with fresh hormone depletion media once an hour, for a total of three hours. Cells were then incubated in hormone depletion media for an additional 24-72 hours prior to induction. Prior to imaging, cells were incubated in fresh hormone depletion media supplemented with 1µM Sir-Hoechst (Cytoskeleton, Cy-SC007). Cells were placed on the microscope in hormone depletion media supplemented with 1nM Sir-Hoechst, and the location of 3-10 regions of interested were recorded. Immediately prior to beginning of time course imaging, 1nM β-estradiol (Millipore-Sigma E8875-250MG) was added directly to directly to imaging media.

#### Nuclear Extraction

Cos7 HaloTag-H2b and Cos7 wild-type cells were grown to approximately 90% confluency in 100mm plates. Cells were collected either by scraping in 4C PBS and centrifugation at 300x G for 3 minutes. The cell pellet was stored at −80°C or immediately processed for nuclear extraction.

For nuclear protein isolation, the cell pellet was resuspended in 500 μL of nuclear extraction buffer containing 10 mM HEPES (Millipore-Sigma, H3375), 10 mM KCl (Millipore-Sigma, P3911), 0.5% Nonidet P-40 Substitute (Millipore-Sigma, 74385), 1 mM DTT (Millipore-Sigma, 11583786001), and 1x cOmplete Mini protease inhibitors (Millipore-Sigma, 11836153001). After a 10-minute room temperature incubation, the sample was centrifuged at 13000xg for 3 minutes to separate the cytoplasmic fraction from the nuclear fraction.

The nuclear pellet was then resuspended in 150 μL of lysis buffer containing 20 mM HEPES pH 8, 400 mM NaCl, 10% glycerol, 1 mM DTT(Millipore-Sigma, 11583786001), 1x cOmplete Mini protease inhibitors (Millipore-Sigma, 11836153001)., and 1 μL/mL Pierce Universal Nuclease (Thermo Fisher Scientific, 88700), and incubated at 4C for 1H. A final centrifugation step at 10000xg for 5 minutes was performed to collect the supernatant containing the nuclear extract. The nuclear extract was then stored at −20C or or used directly for Western blot analysis.

#### Western Blot

Proteins were separated by SDS-PAGE using a 10% polyacrylamide gel and transferred to a PVDF membrane. The membrane was blocked with 3% BSA in TBST for 1 hour at room temperature. Blots were incubated with 500 ng/mL of anti-Histone H2B primary antibody (ABCAM, ab1790) or 1333 ng/mL of Histone H2B Monoclonal Antibody (Bioss, BSM-52099R) overnight at 4°C, followed by incubation with HRP-conjugated secondary antibodies (Thermo Fisher Scientific, 31460) for 1 hour at room temperature. Blots were imaged on a ChemiDoc MP Imaging system (BioRad, 12003154) and quantified using a custom python script.

### Imaging

#### Confocal imaging of Cos7 Halo-H2b cells

Confocal imaging was performed on a Zeiss LSM800 microscope using a 40x Oil immersion objective (NA 1.30) or 63x Water immersion objective (NA 1.20) using standard confocal imaging with a pinhole of 1 AU. Imaging was performed with separate channels for each wavelength, Alexa Fluor 647, Hoechst 33258, and T-PMT and a 2048×2048 pixel field of view.

#### Optical setup

The lattice light sheet imaging system was a modified version of the instrument described in [21]. Key modifications relevant to this work are the use of a greyscale spatial light modulator (Meadowlark P1920-0635-HDMI), a 0.6 NA excitation lens (Thorlabs, TL20X-MPL), and a 1.0 NA detection lens (Zeiss, Objective W “Plan-Apochromat” 20x/1.0, model #421452-9800). To achieve simultaneous multi-color imaging, we split the illumination light into offset strips on the spatial light modulator using a stack of dichroic elements (Semrock Di03-R405, Semrock Di03-R488, Semrock Di03-R561, Semrock MBP01), modulated each wavelength with the indicated lattice pattern on the spatial light modulator, and then recombined the reflected wavefronts by passing back through the same dichroic stack. For the emission path, we used a 642 nm super resolution dichroic (Semrock Di03-R635) to split the emission from JF525 and JF647 into two separate cameras (CamA and CamB respectively). To further filter the emission light and prevent channel bleedthrough from JF525 in For Cos7-HaloTag-H2b and Cos7-HaloTag-NLS imaging, a 514 nm long pass filter (Semrock BLP01-514R) and a 642 nm notch filter (Semrock NF03-642E) was placed in front of CamA. A 647 nm long pass filter (Semrock BLP01-647R) was placed in front of CamB. In addition, a 1000 mm focal length cylindrical lens (Thorlabs LJ516RM) was placed in front of CamB to generate astigmatism for 3D single molecule localization. To compensate for intensity changes in the chromatin channel due to photobleaching, we employed an adaptive power compensation based on the observed channel intensity at each time point. Briefly, after each image stack, we calculated the maximum intensity projection (projected through the z-axis). From this, we identified the 90^th^ percentile pixel value and then linearly increased the laser power based on the change in this value from the previous time point.

Minor alterations in the imaging system were required for imaging of HCT116-mClover-mAID-RPB+Halo-H2b cells labeled with HTL-JF549. A neutral density=1.0 filter was placed in the 560nm excitation path and a notch filter (Semrock NF03-561E) was placed in the emission path in front of CamA rather than 514 nm long pass filter. The optical path for CamB remained the same.

For single molecule localization microscopy (SMLM) we applied a highly inclined swept tile (HIST) microscope similar to the one described in [27]. To axially localize single molecule, a 1000 mm focal length cylindrical lens (Thorlabs LJ516RM) and Semrock FF01-680/42-32 emission filter was placed in front of a Hammamatsu BT Fusion camera.

#### Imaging procedure

For SPT imaging, we imaged each cell over 11 slices with a z-step size of 250 nm. At each slice, we collected 25 consecutive frames. This procedure was repeated for 50 times for each cell. Before and after imaging each cell, the dark current image of both cameras (no excitation was applied) was collected for 1000 frames to estimate the pixel-specific background of the image. For SMLM HIST imaging, we collected 100,000 frames for each field view to have enough number of localizations for reconstruction. The detailed imaging conditions for both cases are listed in the following table.

| Condition | Figure | Illumination mode | Exposure | Laser $\lambda$, power (at objective pupil) |
| --- | --- | --- | --- | --- |
| SPT | Figure1,<br>Figure3,<br>Figure4,<br>Figure S9 | LLS (MB-square, NA 0.4/0.3) | 20 ms | 405 nm, 26 μW |
| | | | | 514 nm, 56 μW |
|  |  |  |  | 642 nm, 9.75 mW |
| SMLM | Figure2,<br>Figure 2s3 | HIST | 25 ms | 642, 40 mW<br>560nm, 11.5 mW |
| MS2 | Figure S8 | LLS (MB-square, NA 0.4/0.3) | 10ms | 642 nm, 25.7 μW<br>488, 27.3 μW |

### Image processing

#### Pre-processing of live cell images

For each cell, a single image for dark current subtraction was generated by taking the mean of the 2000 dark current images that were acquired before and after imaging each cell. The mean dark current image was then subtracted from all dense chromatin and single particle images. Any resulting negative voxel values were set to 0.

To adjust for minor chromatic offsets not corrected by initial microscope alignment, we periodically captured high signal-to-noise ratio volumetric images of densely labeled chromatin in both cameras. We quantified the residual alignment discrepancy between the cameras down to the nearest voxel in all three dimensions (XYZ) using phase cross-correlation. Any detected offsets were then corrected in the dark current-corrected images prior to further image analysis.

#### Densely labeled chromatin

In order to minimize bleed-through of signal from the densely labeled chromatin into simultaneously acquired SPT images, densely labeled chromatin images were acquired under low signal to noise conditions. For each Z position, 25 dark current subtracted images were summed along the t axis and compiled into a high signal-to-noise z-stack. Complied z-stacks were then deskewed to bring into a conventional XYZ reference frame and histogram equalized to the first full z-stack of each cell. To avoid edge artifacts in deconvolution, the deskewed Z-stacks were mirrored across the Z - axis 2 times. Mirrored images were then deconvolved using a Cuda implementation of the Richardson-Lucy algorithm using an experimentally measured LLSM point spread function for 20 iterations (**Movie S6).** The resulting deconvolved images were then cropped to match the dimensions of the original z-stack.

#### SPT and SMLM image processing

Prior to data acquisition, at least 15 z-stacks of 647 nm emission, 100 nm diameter beads were obtained. Super Resolution Microscopy Analysis Platform (SMAP) [73] was used to generate a 3D spline model of our astigmatic point spread function.

SPT images were deskewed to bring them into conventional XYZ reference frame and cropped to match XYZ dimensions as the corresponding deconvolved images. Individual nucleosomes were then localized using SMAP. This was done by identifying regions of the image that contained individual nucleosomes using a difference of Gaussians filter followed by a local maximum filter. The nucleosomes within these regions were then localized in XYZ by fitting the 3D spline model. From these localizations, trajectories were generated using a modified version of uTrack [74]. Our implementation of this algorithm set a maximum linking displacement of 400 nm, a maximum gap length of 2, and disallowed merging and splitting of trajectories. The localizations were converted from nm to voxels through dividing the spatial coordinates of each localization by the voxel size and rounding to the nearest integer value. The voxel values for each localization were then used to determine the chromatin density class, and distance from the edge of the nucleus.

The procedure of HIST SMLM image processing is similar to SPT images. We first generated an astigmatic PSF model and applied it to extract the 3D localization coordinates using SMAP. We used the localization of nanodiamond fiducials to correct for the drift over the course of acquisition. Localizations that occurred within consecutive frames (with a gap of less than two frames) and that were found within a 100 nm radius were linked together into a single localization.

#### Chromatin density classification of live cell images

Nuclear masks for each cell were generated using a custom python script. Briefly, binary images of the chromatin channel were generated using either a Otsu or a multi-Otsu thresholding method. Small holes and small binary objects in the mask there removed using a binary opening, followed by the skimage functions remove_small_holes and remove_small_objects. Following this, all binary objects in the image were indexed using connected components. The largest of these labeled binary objects was kept for subsequent processing as a mask for each individual cell. The specific parameters for these operations, such choice of thresholding method or maximum size of holes, were determined through visual inspection of the resulting masks. These masks were then applied to the deconvolved images prior to the assignment of nuclear of voxels to different chromatin density classes using a previously described maximum likelihood estimation method [25]. Briefly, this approach involves the use of a random field Markov model, which takes into account the relative intensity of the voxel and the chromatin density class of neighboring voxels. After the chromatin density classification, to account for the roll off in intensity at the nuclear edge due to diffraction, all voxels in the least dense chromatin density class (CDC1) that were within 5 voxels of the edge of the nuclear mask were removed from the final chromatin compaction classification image.

#### Scrambled chromatin density classification

Deconvolved images were iteratively rotated and translated in X,Y, and Z. On every i-th iteration, a binary mask of the randomly transformed image was generated, and a binary mask of the i-1 iteration was also created. The voxels of the i-th iteration image that were not already occupied by non-zero values in the i-1 iteration’s image were added to the i-1 image. These iterative transformations resulted in an object that was larger than the original deconvolved nucleus, composed of the randomly transformed gross morphological features of the original nucleus. After the iterative transformations, a binary mask image setting all non-zero voxels in the corresponding chromatin density image equal to one. This binary mask was applied to the randomly transformed image, and then the chromatin density classification was performed (**Figure S3A,B**).

#### Distance from the nuclear periphery

A binary mask was created from the chromatin density class image by assigning a value of one to all non-zero voxels. A distance transform image was generated by calculating the distance of each non-zero voxel to the nearest zero-valued voxel. These distance transform images were used in both determination of the distance of single nucleosomes from the nuclear periphery as well as for voxel-wise comparison of CDCs and distance from the edge of the nucleus.

#### Generation of simulated LLSM images from SMLM localizations

Localizations were grouped into 110 x 110 x 110 nm bins to construct a 3D histogram rendering. Simulated LLSM images of the chromatin were subsequently generated by convolving this 3D histogram with an experimentally measured LLSM point spread function obtained prior to live cell image acquisition. These simulated images were then deconvolved using parameters consistent with those applied in our live cell imaging.

#### Generation of the CDC images from simulated LLSM images and classification of the SMLM localizations

A preliminary binary mask was manually drawn around each cell within the simulated LLSM image. Notably, any intensity regions associated with nanodiamond fiducials were excluded from these masks. For each cell, this mask was applied to the respective image. The highest intensity 110 nm z-plane was identified by summing the intensities along the X and Y axes. Any planes situated more than one z-plane above or below this plane were omitted from further analysis. The ± 1 z-planes were then histogram matched to the central, brightest z plane. The refined single cell LLSM images were then processed and classified into chromatin density classes using the methods previously described.

### Data analysis

#### Analysis of Halo-H2b and Hoechst co-localization

Images of HTL-JF647 and Hoechst were registered using phase cross correlation in order to correct for chromatic aberration. A Gaussian filter with a sigma value of 0.5 was applied to both images. Pixel-wise correlation was then performed using Pearson correlation.

#### Comparison of different number of chromatin density classes

Deconvolved image stacks of densely labeled chromatin images were masked according to the methods described above. The Intensity values for masked, deconvolved images were then flattened into a one-dimensional array and all zero values were dropped. A mixture of gaussian model was fit to the resulting intensity histograms with the number of classes ranging from 1 to 20 and the Akaike information criterion (AIC) and Bayesian information criterion (BIC) were determined for all fits. In order to compare the loss values across cells, the AIC and BIC for fits for all classes were z-score normalized for sample to sample comparison.

#### MSD analysis

MSD was calculated as 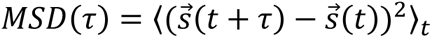. Here 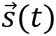 is the localization of a nucleosome at time t in 2D, *τ* is the time interval of the displacement, and 〈*X*〉_*t*_ indicates the average over time. The squared displacements between all localizations within every trajectory were calculated at time intervals of up to 500 ms. The final localization of each displacement was used to assign it to a chromatin density class. MSDs were then averaged across all trajectories of the same class in the same cell, and a linear regression was applied to log(*MSD*) = log(*D*) + *α* ∗ log (*τ*) to extract the diffusion coefficient D and the anomalous exponent α. Extracting anomalous exponent and apparent diffusion coefficient of free diffusing halo tag followed the same method as for nucleosomes.

#### SMLM localization density

For each cell, the number of SMLM localizations was determined for each CDC. The total volume of each CDC was calculated by counting the number of voxels in each CDC in the CDC image and then multiplying by the voxel dimensions in micrometers. Localization density was determined by dividing the total number of localizations in a CDC by its total volume.

#### Power spectrum analysis of live cell and simulated LLSM images

For each cell, a square corresponding to a 5500 nm x 5500 nm in the center of the nucleus was converted from spatial domain to frequency domain using a Fast Fourier Transform. The power at each frequency was computed by calculating the squared magnitude of the Fourier coefficients. A mesh grids representing different frequency components of the image was generated and the overall magnitude of frequency components in the 2D space was calculated. Radial frequencies were binned in bins starting at 0.5 and extending to half the pixel size in increments of 1. For each bin, the average of its boundaries was determined. The squared Fourier amplitudes were then grouped by their mean radial frequency and each binned amplitude was adjusted by multiplying it by the area of the corresponding annular region. The binned amplitudes were normalized by dividing by the maximum binned amplitude.

#### Proportion of voxels in CDCs

For each cell, all CDC images were flattened into a 1D array and all zero values were dropped. From this, a normalized histogram was generated.

#### Visual comparison of SMLM localizations and random localization in a give CDC

The z plane with the highest localization count was identified by scanning a 100nm window in the z dimension and tallying the number of localizations. Localizations outside this z plane were excluded from the super-resolution reconstructions. For each CDC, super-resolution reconstructions were generated by creating a 3D histogram using 20 x 20 x 100 nm (XYZ) bins. Lines indicating the voxel boundaries of in the CDC image were drawn manually on the CDC image and then overlaid on the reconstructed SMLM image. For visual comparison, we generated randomly distributed localizations of the same density as our experimental measurements and plotted them within same CDC boundaries (**Figure S5**).

#### Analysis of voxel wise chromatin intensity/ CDC dynamics

A 1 x 6 voxel line profile was arbitrarily chosen in an example cell. The values for each voxel in the line profile were then recorded at all time points obtained (**Figure S1E**).

#### Radius of gyration analysis

For each trajectory in a given cell, all localizations were registered to the origin by subtracting the mean XY location of the trajectory from all localizations in said trajectory. The registered localizations were then segmented according the CDC of the original localization. The radius of gyration was determined for each CDC by taking the root mean squared distance of from the origin.

#### Angular anisotropy analysis

The angle between every set of three subsequent localizations in a trajectory was determined. Said angles were then partitioned according to the CDC of the middle of the three points. Displacements were classified as ‘forward’ if the angle between displacements was between −30 and 30 degrees and ‘backward’ if the angle between displacements was between 150 and 210 degrees. For each CDC, the fold change between forward and backward displacements was determined by taking the log_2_ of the number of forward displacements divided by the number of backward displacements.

#### Pair correlation function calculation and fractal dimension estimation

The normalized pair correlation function (G(r)) was calculated in a similar approach as described in [75], where 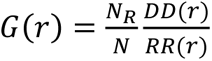. Here N is the total number of localizations in the actual data, and N_R_ is the total number of localizations randomly distributed with the same density as the actual data. DD(r) is the average number of pairs of objects with separation r in the actual data, and RR(r) is the average number of pairs of objects with separation r in the randomly distributed data. To account for diminished localization detection efficiency away from the focal plane, we adopted an approached similar to [33]. Briefly, for each cell, we plotted a histogram of the number of localizations found at each z-plane. We fit this histogram to a Gaussian function and then scaled the number of randomly distributed points in 3D space by this same Gaussian function. In this manner, the spatially random points were subject to the same axial sampling bias as our experimental measurements.

Based on the G(r) curve, we performed a linear regression on to log(G(r)) and log(r) to extract the decaying exponent γ of G(r). The lower bound of the linear regression was set at 35 nm. The upper bound of the linear regression was determined through iteratively increasing the maximum value until the r^2^ value of the fit fell below 0.98 ranging from the 35 nm to an upper value that was determined by a r^2^ value of 0.98. The slope of this regression represents γ and decays approximately as *G*(*r*) ∝ *r*^−γ^. The fractal dimension is calculated as *d*_*f*_ = 3 − γ.

#### Whole cell G(r) analysis

For each r value in our G(r) plot, spanning from 75 nm to 565 nm, we fit two linear regressions. The first regression used values from 75 nm to the r value, while the second regression covered values from the r value up to 505 nm. We then calculated the summed squared residual for both regressions across the respective ranges. The optimal hinge point was selected based on the r value that produced the smallest combined summed squared residual. The estimated fractal dimension was derived from the slopes of these regressions, consistent with previously detailed methods.

#### Calculation of model predicted MSD characterizations

Our overall procedure follows the formulas described in [39]. Briefly, the predicted anomalous exponent for nucleosomes is calculated as 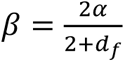, where *α* is the anomalous exponent of free diffusing halotag and *d*_*f*_ is the chromatin fractal dimension. The predicted nucleosome apparent diffusion coefficient is calculated as 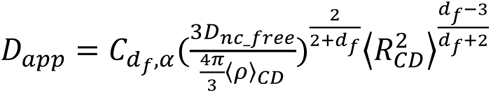, where 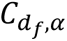 is a constant that depends on *d*_*f*_ and *α* and follows the expression 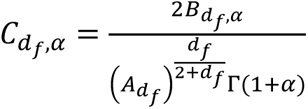, with 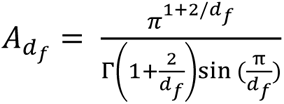, and 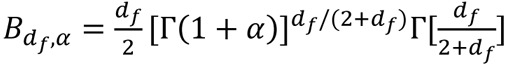. This expression is based on equation 15 and 17 in [39] and we substitute 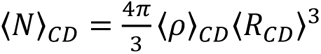. **Dnc*_*free** is the diffusion coefficient of freely diffusing (non-chromatin bound) nucleosomes. This is different from the experimentally measured nucleosome diffusion coefficient as they are connected by DNA. We estimated **Dnc*_*free** by scaling the measured diffusion coefficient of halotag by the relative hydrodynamic radii of halotag and nucleosomes, 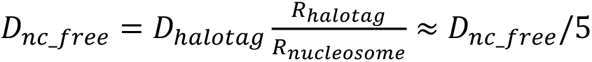. ⟨*ρ*⟩*_CD_* is the density of nucleosomes in chromatin domain, which we estimated based on the localization density in the super-resolution nucleosome data, and corrected for the blinking characterization of JF630B dye and the labelling efficiency of halo-tag-H2b (described in more detail in the next section, **Figure S1B**). 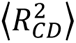 is the averaged square size of the chromatin domain, and we estimated it based on the length where the slope of pair correlation function reaches zero (**Figure S6D and E**). The errors in the predicted anomalous exponent *β* and the predicted apparent diffusion coefficient of nucleosomes *D*_*app*_ are estimated by propagating the errors in *α* and *d*_*f*_ as estimated from experimental measurements.

#### Determination of nucleosome density

Estimated nucleosome density is calculated as 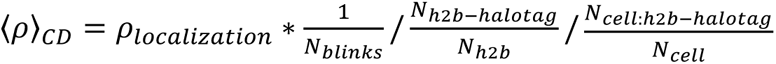, where **ρ*_*localization*_* is the density of localization, *N_*blinks*_* is the average number of blinks of each dye molecule determined by **Figure S6 A, B**, 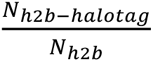 is the ratio of Halotag-H2b to endogenous h2b, which is determined by the Western blot in **Figure S1B**, and 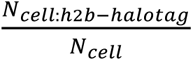 is the fraction of cells that express h2b-halo-tag determined by **Figure S6C**.

## References

[1] F. Grasser, M. Neusser, H. Fiegler, T. Thormeyer, M. Cremer, N.P. Carter, T. Cremer, S. Müller, Replication-timing-correlated spatial chromatin arrangements in cancer and in primate interphase nuclei, J. Cell Sci. 121 (2008) 1876–1886. 10.1242/jcs.026989.

[2] Y. Chen, Y. Zhang, Y. Wang, L. Zhang, E.K. Brinkman, S.A. Adam, R. Goldman, B. van Steensel, J. Ma, A.S. Belmont, Mapping 3D genome organization relative to nuclear compartments using TSA-Seq as a cytological rulerTSA-Seq mapping of nuclear genome organization, J. Cell Biol. 217 (2018) 4025–4048. 10.1083/jcb.201807108.

[3] S.A. Quinodoz, N. Ollikainen, B. Tabak, A. Palla, J.M. Schmidt, E. Detmar, M.M. Lai, A.A. Shishkin, P. Bhat, Y. Takei, V. Trinh, E. Aznauryan, P. Russell, C. Cheng, M. Jovanovic, A. Chow, L. Cai, P. McDonel, M. Garber, M. Guttman, Higher-Order Inter-chromosomal Hubs Shape 3D Genome Organization in the Nucleus, Cell. 174 (2018) 744–757.e24. 10.1016/j.cell.2018.05.024.

[4] D.L. Spector, W.H. Schrier, H. Busch, Immunoelectron microscopic localization of snRNPs, Biol. Cell. 49 (1984) 1–10. 10.1111/j.1768-322X.1984.tb00215.x.

[5] D.D. Brown, J.B. Gurdon, Absence of ribosomal rna synthesis in the anucleolate mutant of xenopus laevis, Proc. Natl. Acad. Sci. 51 (1964) 139–146. 10.1073/pnas.51.1.139.

[6] R.D. Kornberg, Chromatin Structure: A Repeating Unit of Histones and DNA, Science. 184 (1974) 868–871.

[7] H.D. Ou, S. Phan, T.J. Deerinck, A. Thor, M.H. Ellisman, C.C. O’Shea, ChromEMT: Visualizing 3D chromatin structure and compaction in interphase and mitotic cells, Science. 357 (2017) eaag0025. 10.1126/science.aag0025.

[8] M.A. Ricci, C. Manzo, M.F. García-Parajo, M. Lakadamyali, M.P. Cosma, Chromatin Fibers Are Formed by Heterogeneous Groups of Nucleosomes In Vivo, Cell. 160 (2015) 1145– 1158. 10.1016/j.cell.2015.01.054.

[9] E. Miron, R. Oldenkamp, J.M. Brown, D.M.S. Pinto, C.S. Xu, A.R. Faria, H.A. Shaban, J.D.P. Rhodes, C. Innocent, S. de Ornellas, H.F. Hess, V. Buckle, L. Schermelleh, Chromatin arranges in chains of mesoscale domains with nanoscale functional topography independent of cohesin, Sci. Adv. 6 (2020) eaba8811. 10.1126/sciadv.aba8811.

[10] T. Nozaki, R. Imai, M. Tanbo, R. Nagashima, S. Tamura, T. Tani, Y. Joti, M. Tomita, K. Hibino, M.T. Kanemaki, K.S. Wendt, Y. Okada, T. Nagai, K. Maeshima, Dynamic Organization of Chromatin Domains Revealed by Super-Resolution Live-Cell Imaging, Mol. Cell. 67 (2017) 282–293.e7. 10.1016/j.molcel.2017.06.018.

[11] S.M. Stack, D.B. Brown, W.C. Dewey, Visualization of interphase chromosomes, J. Cell Sci. 26 (1977) 281–299. 10.1242/jcs.26.1.281.

[12] C. Zorn, T. Cremer, C. Cremer, J. Zimmer, Laser UV microirradiation of interphase nuclei and post-treatment with caffeine, Hum. Genet. 35 (1976) 83–89. 10.1007/BF00295622.

[13] L.A. Mirny, The fractal globule as a model of chromatin architecture in the cell, Chromosome Res. 19 (2011) 37–51. 10.1007/s10577-010-9177-0.

[14] T. Nozaki, S. Shinkai, S. Ide, K. Higashi, S. Tamura, M.A. Shimazoe, M. Nakagawa, Y. Suzuki, Y. Okada, M. Sasai, S. Onami, K. Kurokawa, S. Iida, K. Maeshima, Condensed but liquid-like domain organization of active chromatin regions in living human cells, Sci. Adv. 9 (2023) eadf1488. 10.1126/sciadv.adf1488.

[15] P.A. Gómez-García, S. Portillo-Ledesma, M.V. Neguembor, M. Pesaresi, W. Oweis, T. Rohrlich, S. Wieser, E. Meshorer, T. Schlick, M.P. Cosma, M. Lakadamyali, Mesoscale Modeling and Single-Nucleosome Tracking Reveal Remodeling of Clutch Folding and Dynamics in Stem Cell Differentiation, Cell Rep. 34 (2021) 108614. 10.1016/j.celrep.2020.108614.

[16] M. Cremer, K. Brandstetter, A. Maiser, S.S.P. Rao, V.J. Schmid, M. Guirao-Ortiz, N. Mitra, S. Mamberti, K.N. Klein, D.M. Gilbert, H. Leonhardt, M.C. Cardoso, E.L. Aiden, H. Harz, T. Cremer, Cohesin depleted cells rebuild functional nuclear compartments after endomitosis, Nat. Commun. 11 (2020) 6146. 10.1038/s41467-020-19876-6.

[17] R. Nagashima, K. Hibino, S.S. Ashwin, M. Babokhov, S. Fujishiro, R. Imai, T. Nozaki, S. Tamura, T. Tani, H. Kimura, M. Shribak, M.T. Kanemaki, M. Sasai, K. Maeshima, Single nucleosome imaging reveals loose genome chromatin networks via active RNA polymerase II, J. Cell Biol. 218 (2019) 1511–1530. 10.1083/jcb.201811090.

[18] M.N. Saxton, T. Morisaki, D. Krapf, H. Kimura, T.J. Stasevich, Live-cell imaging uncovers the relationship between histone acetylation, transcription initiation, and nucleosome mobility, (2023) 2023.03.02.530854. 10.1101/2023.03.02.530854.

[19] S. Iida, S. Shinkai, Y. Itoh, S. Tamura, M.T. Kanemaki, S. Onami, K. Maeshima, Single-nucleosome imaging reveals steady-state motion of interphase chromatin in living human cells, Sci. Adv. 8 (2022) eabn5626. 10.1126/sciadv.abn5626.

[20] A. Bancaud, S. Huet, N. Daigle, J. Mozziconacci, J. Beaudouin, J. Ellenberg, Molecular crowding affects diffusion and binding of nuclear proteins in heterochromatin and reveals the fractal organization of chromatin, EMBO J. 28 (2009) 3785–3798. 10.1038/emboj.2009.340.

[21] B.-C. Chen, W.R. Legant, K. Wang, L. Shao, D.E. Milkie, M.W. Davidson, C. Janetopoulos, X.S. Wu, J.A. Hammer, Z. Liu, B.P. English, Y. Mimori-Kiyosue, D.P. Romero, A.T. Ritter, J. Lippincott-Schwartz, L. Fritz-Laylin, R.D. Mullins, D.M. Mitchell, J.N. Bembenek, A.-C. Reymann, R. Böhme, S.W. Grill, J.T. Wang, G. Seydoux, U.S. Tulu, D.P. Kiehart, E. Betzig, Lattice light-sheet microscopy: Imaging molecules to embryos at high spatiotemporal resolution, Science. 346 (2014) 1257998. 10.1126/science.1257998.

[22] Y. Shi, T.A. Daugird, W.R. Legant, A quantitative analysis of various patterns applied in lattice light sheet microscopy, Nat. Commun. 13 (2022) 4607. 10.1038/s41467-022-32341-w.

[23] J.B. Grimm, B.P. English, J. Chen, J.P. Slaughter, Z. Zhang, A. Revyakin, R. Patel, J.J. Macklin, D. Normanno, R.H. Singer, T. Lionnet, L.D. Lavis, A general method to improve fluorophores for live-cell and single-molecule microscopy, Nat. Methods. 12 (2015) 244–250. 10.1038/nmeth.3256.

[24] J.-H. Su, P. Zheng, S.S. Kinrot, B. Bintu, X. Zhuang, Genome-Scale Imaging of the 3D Organization and Transcriptional Activity of Chromatin, Cell. 182 (2020) 1641–1659.e26. 10.1016/j.cell.2020.07.032.

[25] V.J. Schmid, M. Cremer, T. Cremer, Quantitative analyses of the 3D nuclear landscape recorded with super-resolved fluorescence microscopy, Methods San Diego Calif. 123 (2017) 33–46. 10.1016/j.ymeth.2017.03.013.

[26] C. Deo, A.S. Abdelfattah, H.K. Bhargava, A.J. Berro, N. Falco, H. Farrants, B. Moeyaert, M. Chupanova, L.D. Lavis, E.R. Schreiter, The HaloTag as a general scaffold for far-red tunable chemigenetic indicators, Nat. Chem. Biol. 17 (2021) 718–723. 10.1038/s41589-021-00775-w.

[27] J. Tang, K.Y. Han, Extended field-of-view single-molecule imaging by highly inclined swept illumination, Optica. 5 (2018) 1063–1069. 10.1364/OPTICA.5.001063.

[28] P.J.E. Peebles, Statistical Analysis of Catalogs of Extragalactic Objects. I. Theory, Astrophys. J. 185 (1973) 413. 10.1086/152431.

[29] M. Ester, H.-P. Kriegel, J. Sander, X. Xu, A density-based algorithm for discovering clusters in large spatial databases with noise, in: Proc. Second Int. Conf. Knowl. Discov. Data Min., AAAI Press, Portland, Oregon, 1996: pp. 226–231.

[30] E. Schubert, J. Sander, M. Ester, H.P. Kriegel, X. Xu, DBSCAN Revisited, Revisited: Why and How You Should (Still) Use DBSCAN, ACM Trans. Database Syst. 42 (2017) 19:1–19:21. 10.1145/3068335.

[31] F. Levet, E. Hosy, A. Kechkar, C. Butler, A. Beghin, D. Choquet, J.-B. Sibarita, SR-Tesseler: a method to segment and quantify localization-based super-resolution microscopy data, Nat. Methods. 12 (2015) 1065–1071. 10.1038/nmeth.3579.

[32] G. Voronoi, Nouvelles applications des paramètres continus à la théorie des formes quadratiques. Premier mémoire. Sur quelques propriétés des formes quadratiques positives parfaites., J. Für Reine Angew. Math. Crelles J. 1908 (1908) 97–102. 10.1515/crll.1908.133.97.

[33] V. Récamier, I. Izeddin, L. Bosanac, M. Dahan, F. Proux, X. Darzacq, Single cell correlation fractal dimension of chromatin, Nucleus. 5 (2014) 75–84. 10.4161/nucl.28227.

[34] A. Bancaud, C. Lavelle, S. Huet, J. Ellenberg, A fractal model for nuclear organization: current evidence and biological implications, Nucleic Acids Res. 40 (2012) 8783–8792. 10.1093/nar/gks586.

[35] E. Lieberman-Aiden, N.L. van Berkum, L. Williams, M. Imakaev, T. Ragoczy, A. Telling, I. Amit, B.R. Lajoie, P.J. Sabo, M.O. Dorschner, R. Sandstrom, B. Bernstein, M.A. Bender, M. Groudine, A. Gnirke, J. Stamatoyannopoulos, L.A. Mirny, E.S. Lander, J. Dekker, Comprehensive Mapping of Long-Range Interactions Reveals Folding Principles of the Human Genome, Science. 326 (2009) 289–293. 10.1126/science.1181369.

[36] R.K. Sachs, G. van den Engh, B. Trask, H. Yokota, J.E. Hearst, A random-walk/giant-loop model for interphase chromosomes., Proc. Natl. Acad. Sci. 92 (1995) 2710–2714. 10.1073/pnas.92.7.2710.

[37] P.J. Fleming, K.G. Fleming, HullRad: Fast Calculations of Folded and Disordered Protein and Nucleic Acid Hydrodynamic Properties, Biophys. J. 114 (2018) 856–869. 10.1016/j.bpj.2018.01.002.

[38] K.E. Polovnikov, B. Slavov, S. Belan, M. Imakaev, H.B. Brandão, L.A. Mirny, Crumpled polymer with loops recapitulates key features of chromosome organization, (2023) 2022.02.01.478588. 10.1101/2022.02.01.478588.

[39] S. Shinkai, T. Nozaki, K. Maeshima, Y. Togashi, Dynamic Nucleosome Movement Provides Structural Information of Topological Chromatin Domains in Living Human Cells, PLOS Comput. Biol. 12 (2016) e1005136. 10.1371/journal.pcbi.1005136.

[40] A. Yesbolatova, Y. Saito, N. Kitamoto, H. Makino-Itou, R. Ajima, R. Nakano, H. Nakaoka, K. Fukui, K. Gamo, Y. Tominari, H. Takeuchi, Y. Saga, K. Hayashi, M.T. Kanemaki, The auxin-inducible degron 2 technology provides sharp degradation control in yeast, mammalian cells, and mice, Nat. Commun. 11 (2020) 5701. 10.1038/s41467-020-19532-z.

[41] E.C. Cesconetto, F.S.A. Junior, F. a. P. Crisafuli, O.N. Mesquita, E.B. Ramos, M.S. Rocha, DNA interaction with Actinomycin D: mechanical measurements reveal the details of the binding data, Phys. Chem. Chem. Phys. PCCP. 15 (2013) 11070–11077. 10.1039/c3cp50898f.

[42] J. Otterstrom, A. Castells-Garcia, C. Vicario, P.A. Gomez-Garcia, M.P. Cosma, M. Lakadamyali, Super-resolution microscopy reveals how histone tail acetylation affects DNA compaction within nucleosomes in vivo, Nucleic Acids Res. 47 (2019) 8470–8484. 10.1093/nar/gkz593.

[43] K.F. Tóth, T.A. Knoch, M. Wachsmuth, M. Frank-Stöhr, M. Stöhr, C.P. Bacher, G. Müller, K. Rippe, Trichostatin A-induced histone acetylation causes decondensation of interphase chromatin, J. Cell Sci. 117 (2004) 4277–4287. 10.1242/jcs.01293.

[44] J. Rodriguez, G. Ren, C.R. Day, K. Zhao, C.C. Chow, D.R. Larson, Intrinsic Dynamics of a Human Gene Reveal the Basis of Expression Heterogeneity, Cell. 176 (2019) 213–226.e18. 10.1016/j.cell.2018.11.026.

[45] K.M. Creamer, H.J. Kolpa, J.B. Lawrence, Nascent RNA scaffolds contribute to chromosome territory architecture and counter chromatin compaction, Mol. Cell. 81 (2021) 3509–3525.e5. 10.1016/j.molcel.2021.07.004.

[46] J. Lou, L. Scipioni, B.K. Wright, T.K. Bartolec, J. Zhang, V.P. Masamsetti, K. Gaus, E. Gratton, A.J. Cesare, E. Hinde, Phasor histone FLIM-FRET microscopy quantifies spatiotemporal rearrangement of chromatin architecture during the DNA damage response, Proc. Natl. Acad. Sci. U. S. A. 116 (2019) 7323–7332. 10.1073/pnas.1814965116.

[47] M.V. Neguembor, L. Martin, Á. Castells-García, P.A. Gómez-García, C. Vicario, D. Carnevali, J.A. Abed, A. Granados, R. Sebastian-Perez, F. Sottile, J. Solon, C. Wu, M. Lakadamyali, M.P. Cosma, Transcription-mediated supercoiling regulates genome folding and loop formation, Mol. Cell. 81 (2021) 3065–3081.e12. 10.1016/j.molcel.2021.06.009.

[48] M.B. Felisbino, M.S.V. Gatti, M.L.S. Mello, Changes in Chromatin Structure in NIH 3T3 Cells Induced by Valproic Acid and Trichostatin A, J. Cell. Biochem. 115 (2014) 1937– 1947. 10.1002/jcb.24865.

[49] P. Bhat, D. Honson, M. Guttman, Nuclear compartmentalization as a mechanism of quantitative control of gene expression, Nat. Rev. Mol. Cell Biol. 22 (2021) 653–670. 10.1038/s41580-021-00387-1.

[50] H.A. Shaban, R. Barth, L. Recoules, K. Bystricky, Hi-D: nanoscale mapping of nuclear dynamics in single living cells, Genome Biol. 21 (2020) 95. 10.1186/s13059-020-02002-6.

[51] S. Grosse-Holz, A. Coulon, L. Mirny, Scale-free models of chromosome structure, dynamics, and mechanics, (2023) 2023.04.14.536939. 10.1101/2023.04.14.536939.

[52] V.I.P. Keizer, S. Grosse-Holz, M. Woringer, L. Zambon, K. Aizel, M. Bongaerts, F. Delille, L. Kolar-Znika, V.F. Scolari, S. Hoffmann, E.J. Banigan, L.A. Mirny, M. Dahan, D. Fachinetti, A. Coulon, Live-cell micromanipulation of a genomic locus reveals interphase chromatin mechanics, Science. 377 (2022) 489–495. 10.1126/science.abi9810.

[53] B. van Steensel, A.S. Belmont, Lamina-Associated Domains: Links with Chromosome Architecture, Heterochromatin, and Gene Repression, Cell. 169 (2017) 780–791. 10.1016/j.cell.2017.04.022.

[54] A. Gonzalez-Sandoval, B.D. Towbin, V. Kalck, D.S. Cabianca, D. Gaidatzis, M.H. Hauer, L. Geng, L. Wang, T. Yang, X. Wang, K. Zhao, S.M. Gasser, Perinuclear Anchoring of H3K9-Methylated Chromatin Stabilizes Induced Cell Fate in C. elegans Embryos, Cell. 163 (2015) 1333–1347. 10.1016/j.cell.2015.10.066.

[55] A. Poleshko, K.M. Mansfield, C.C. Burlingame, M.D. Andrake, N.R. Shah, R.A. Katz, The Human Protein PRR14 Tethers Heterochromatin to the Nuclear Lamina during Interphase and Mitotic Exit, Cell Rep. 5 (2013) 292–301. 10.1016/j.celrep.2013.09.024.

[56] L. Guelen, L. Pagie, E. Brasset, W. Meuleman, M.B. Faza, W. Talhout, B.H. Eussen, A. de Klein, L. Wessels, W. de Laat, B. van Steensel, Domain organization of human chromosomes revealed by mapping of nuclear lamina interactions, Nature. 453 (2008) 948–951. 10.1038/nature06947.

[57] D. Peric-Hupkes, W. Meuleman, L. Pagie, S.W.M. Bruggeman, I. Solovei, W. Brugman, S. Gräf, P. Flicek, R.M. Kerkhoven, M. van Lohuizen, M. Reinders, L. Wessels, B. van Steensel, Molecular Maps of the Reorganization of Genome-Nuclear Lamina Interactions during Differentiation, Mol. Cell. 38 (2010) 603–613. 10.1016/j.molcel.2010.03.016.

[58] B. Wen, H. Wu, Y. Shinkai, R.A. Irizarry, A.P. Feinberg, Large histone H3 lysine 9 dimethylated chromatin blocks distinguish differentiated from embryonic stem cells, Nat. Genet. 41 (2009) 246–250. 10.1038/ng.297.

[59] Y. Li, A. Eshein, R.K.A. Virk, A. Eid, W. Wu, J. Frederick, D. VanDerway, S. Gladstein, K. Huang, A.R. Shim, N.M. Anthony, G.M. Bauer, X. Zhou, V. Agrawal, E.M. Pujadas, S. Jain, G. Esteve, J.E. Chandler, T.-Q. Nguyen, R. Bleher, J.J. de Pablo, I. Szleifer, V.P. Dravid, L.M. Almassalha, V. Backman, Nanoscale chromatin imaging and analysis platform bridges 4D chromatin organization with molecular function, Sci. Adv. 7 (2021) eabe4310. 10.1126/sciadv.abe4310.

[60] K. Luger, A.W. Mäder, R.K. Richmond, D.F. Sargent, T.J. Richmond, Crystal structure of the nucleosome core particle at 2.8 Å resolution, Nature. 389 (1997) 251–260. 10.1038/38444.

[61] S.M. Görisch, M. Wachsmuth, K.F. Tóth, P. Lichter, K. Rippe, Histone acetylation increases chromatin accessibility, J. Cell Sci. 118 (2005) 5825–5834. 10.1242/jcs.02689.

[62] S.C. Weber, A.J. Spakowitz, J.A. Theriot, Bacterial Chromosomal Loci Move Subdiffusively through a Viscoelastic Cytoplasm, Phys. Rev. Lett. 104 (2010) 238102. 10.1103/PhysRevLett.104.238102.

[63] G.L. Hunter, E.R. Weeks, The physics of the colloidal glass transition, Rep. Prog. Phys. 75 (2012) 066501. 10.1088/0034-4885/75/6/066501.

[64] G. Shi, L. Liu, C. Hyeon, D. Thirumalai, Interphase human chromosome exhibits out of equilibrium glassy dynamics, Nat. Commun. 9 (2018) 3161. 10.1038/s41467-018-05606-6.

[65] H. Ohishi, S. Shimada, S. Uchino, J. Li, Y. Sato, M. Shintani, H. Owada, Y. Ohkawa, A. Pertsinidis, T. Yamamoto, H. Kimura, H. Ochiai, STREAMING-tag system reveals spatiotemporal relationships between transcriptional regulatory factors and transcriptional activity, Nat. Commun. 13 (2022) 7672. 10.1038/s41467-022-35286-2.

[66] B. Gu, T. Swigut, A. Spencley, M.R. Bauer, M. Chung, T. Meyer, J. Wysocka, Transcription-coupled changes in nuclear mobility of mammalian cis-regulatory elements, Science. 359 (2018) 1050–1055. 10.1126/science.aao3136.

[67] Y. rhie Y. Xu, X. Chen, S. Feng, Z. Liu, Y. Sun, X. Yao, F. Li, W. Zhu, L. Gao, H. Chen, Z. Du, W. Xie, X. Xu, X. Huang, J. Liu, 3D Chromatin Structures of Mature Gametes and Structural Reprogramming during Mammalian Embryogenesis, Cell. 170 (2017) 367–381.e20. 10.1016/j.cell.2017.06.029.

[68] S.K. Rhie, A.A. Perez, F.D. Lay, S. Schreiner, J. Shi, J. Polin, P.J. Farnham, A high-resolution 3D epigenomic map reveals insights into the creation of the prostate cancer transcriptome, Nat. Commun. 10 (2019) 4154. 10.1038/s41467-019-12079-8.

[69] J. Xu, H. Ma, H. Ma, W. Jiang, C.A. Mela, M. Duan, S. Zhao, C. Gao, E.-R. Hahm, S.M. Lardo, K. Troy, M. Sun, R. Pai, D.B. Stolz, L. Zhang, S. Singh, R.E. Brand, D.J. Hartman, J. Hu, S.J. Hainer, Y. Liu, Super-resolution imaging reveals the evolution of higher-order chromatin folding in early carcinogenesis, Nat. Commun. 11 (2020) 1899. 10.1038/s41467-020-15718-7.

[70] J.R. Dixon, I. Jung, S. Selvaraj, Y. Shen, J.E. Antosiewicz-Bourget, A.Y. Lee, Z. Ye, A. Kim, N. Rajagopal, W. Xie, Y. Diao, J. Liang, H. Zhao, V.V. Lobanenkov, J.R. Ecker, J.A. Thomson, B. Ren, Chromatin architecture reorganization during stem cell differentiation, Nature. 518 (2015) 331–336. 10.1038/nature14222.

[71] L. Li, H. Liu, P. Dong, D. Li, W.R. Legant, J.B. Grimm, L.D. Lavis, E. Betzig, R. Tjian, Z. Liu, Real-time imaging of Huntingtin aggregates diverting target search and gene transcription, eLife. 5 (2016) e17056. 10.7554/eLife.17056.

[72] N.C. Yeo, A. Chavez, A. Lance-Byrne, Y. Chan, D. Menn, D. Milanova, C.-C. Kuo, X. Guo, S. Sharma, A. Tung, R.J. Cecchi, M. Tuttle, S. Pradhan, E.T. Lim, N. Davidsohn, M.R. Ebrahimkhani, J.J. Collins, N.E. Lewis, S. Kiani, G.M. Church, An enhanced CRISPR repressor for targeted mammalian gene regulation, Nat. Methods. 15 (2018) 611–616. 10.1038/s41592-018-0048-5.

[73] J. Ries, SMAP: a modular super-resolution microscopy analysis platform for SMLM data, Nat. Methods. 17 (2020) 870–872. 10.1038/s41592-020-0938-1.

[74] K. Jaqaman, D. Loerke, M. Mettlen, H. Kuwata, S. Grinstein, S.L. Schmid, G. Danuser, Robust single-particle tracking in live-cell time-lapse sequences, Nat. Methods. 5 (2008) 695–702. 10.1038/nmeth.1237.

[75] L. Xie, P. Dong, X. Chen, T.-H.S. Hsieh, S. Banala, M. De Marzio, B.P. English, Y. Qi, S.K. Jung, K.-R. Kieffer-Kwon, W.R. Legant, A.S. Hansen, A. Schulmann, R. Casellas, B. Zhang, E. Betzig, L.D. Lavis, H.Y. Chang, R. Tjian, Z. Liu, 3D ATAC-PALM: super-resolution imaging of the accessible genome, Nat. Methods. 17 (2020) 430–436. 10.1038/s41592-020-0775-2.

