## Supplemental Material for "Correlative single molecule lattice light sheet imaging reveals the dynamic relationship between nucleosomes and the local chromatin environment"

### Supplemental Materials

#### Table of Contents


List of SI movies 2
Supplementary figures 3-12

**List of SI movies**

**Movie S1:** 3D lattice light sheet microscopy and classification of chromatin density in live cells.

**Movie S2:** Correlative single molecule tracking and lattice light sheet microscopy of nucleosomes and chromatin.

**Movie S3:** 3D highly inclined and swept tile imaging and classification of nucleosome organization.

**Movie S4:** Correlative single molecule tracking and lattice light sheet microscopy of free diffusing Halotag-NLS and chromatin.

**Movie S5:** Multicolor lattice light sheet microscopy of transcriptional bursts and chromatin.

**Movie S6: P**rocessing steps chromatin images from lattice light sheet microscope.

#### Supplementary Figures and Captions:

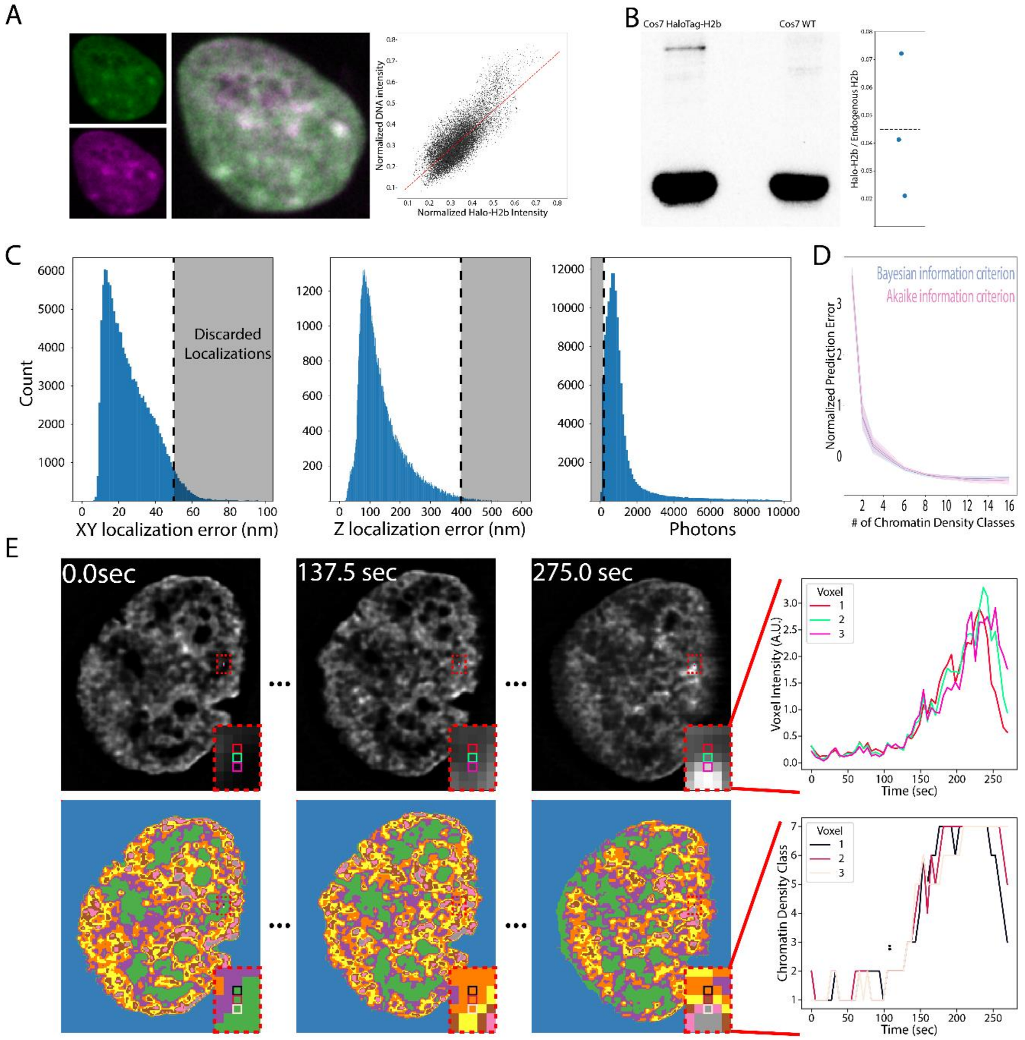


**Figure S1: (A)** Confocal images of DNA (Green, Hoechst) and HaloTag-H2b (Magenta, HTL-JF647) in Cos7-HaloTag-H2b cells. The middle panel shows the merged image. The right panel shows the pixel-wise intensity correlation between the two. The red line shows the linear fit of the data, with $R^2 = 0.77$. **(B)** Example western blot to quantify the ratio of HaloTag-H2b to endogenous H2b in Cos7-HaloTag-H2b cells and Cos7 wildtype cells. Quantification of the exogenous HaloTag-H2b band relative to endogenous H2b band. Blue dots represent one replicate and dashed line represent the mean of all replicates. **(C)** The distribution of nucleosomes' lateral (left) and axial (middle) localization error and the number of photons (right) for live-cell lattice light sheet imaging. Shaded areas are discarded by the post-processing localization filter. **(D)** Normalized Akaike information criterion (magenta) and Bayesian information criterion (blue) for fitting of mixture of gaussian model with increasing number of components to intensity histograms of deconvolved chromatin images. Values are z-score normalized. Solid lines represent the mean value and the shaded regions represent the 95% confidence interval. **(E)** Top panel: Representative deconvolved chromatin images taken at the beginning, half way through and at end of imaging. Line plot indicating the fluorescence intensity (A.U) as a function of time (seconds) of set of voxels in a 3x1 line profile drawn in the deconvolved image. Bottom panel: Representative chromatin density classification images of top panel. Line plot indicating the chromatin density class as a function of time (seconds) of for the same set of voxels in top panel. Data in A are from a single representative cell. Data in B are from n = 3 independent biological replicates. Data in C are representative distributions from a single cell. Data in D are from n = 20 cells from 1 biological replicate.

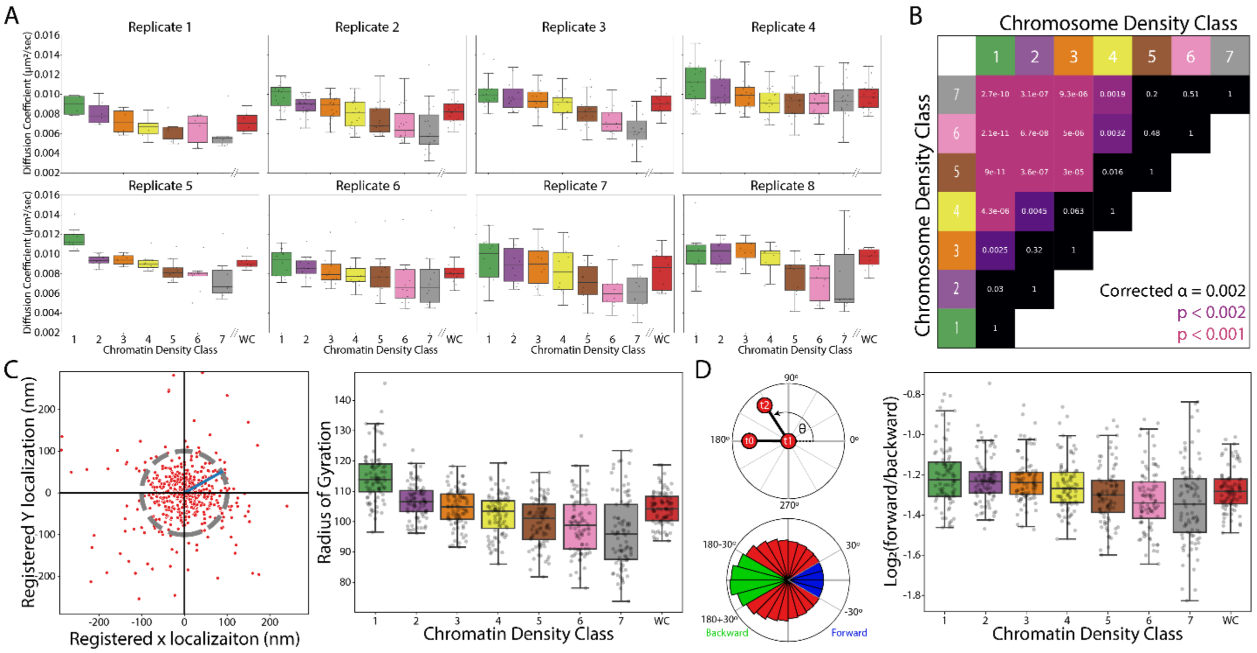


**Figure S2: (A)** Box plot of the apparent diffusion coefficient of nucleosome motion in different chromatin classes across multiple experimental replicates. The figure convention is the same as in Figure1E. WC indicates whole cell. **(B)** Pair-wise t-test of nucleosome diffusion coefficient between different chromatin classes. **(C)** Left: Schematics illustrating the calculation of the radius of gyration. Red circles represent individual localizations, registered to the origin. The derived radius of gyration is depicted as a blue line, which in turn defines the gray circle centered around the origin. Right: box plot of radius of gyration in different chromatin density classes, figure follows the same convention as Figure1G. **(D)** Left: schematics for calculating the anisotropy of nucleosome motion. θ is defined as the angle between two consecutive steps. For θ between -30 to 30 degrees, the motion is defined as forward; for θ between 150 to 210 degrees, the motion is defined as backward. Right: box plot of the ratio of fold change of forward/backward portion in different chromatin classes. Figure follows the same convention as Figure1G. Data in A and B are from n = 88 cells across 8 independent biological replicates. Data in C and D are from n = 88 cells across 8 independent biological replicates.

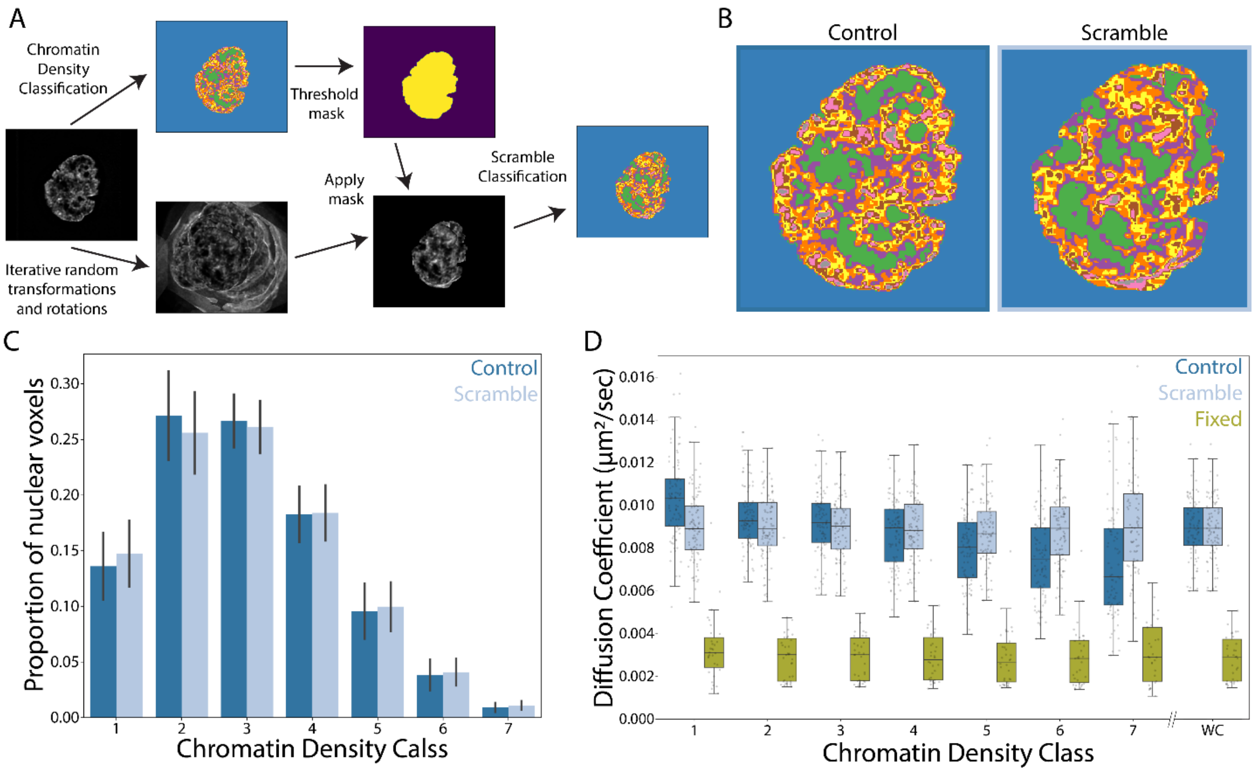


**Figure S3: (A)** Schematic for generating scrambled chromatin classes. A scrambled, space filling image is generated through iteratively adding a transformed and rotated chromatin image. This randomly transformed image is masked, and chromatin density classification is performed. **(B)** Comparison between the real chromatin density classes (left) and the scrambled chromatin density classes (right). **(C)** Histogram of nuclear voxels associated chromatin density classes for chromatin (dark blue) and scrambled (light blue) conditions. Bar height indicates the mean proportion of voxels in a given class. The error bars indicate standard deviation. **(D)** Box plot of diffusion coefficient in control (dark blue, same as Figure 1G) and scrambled chromatin (light blue) classes. The yellow boxes show the diffusion coefficient of nucleosomes in chemically fixed cells. Data in C and D are from n = 88 cells across 8 independent biological replicates (Control), n = 88 cells across 8 independent biological replicates (Scramble), n = 34 cells across 3 independent biological replicates (Fixed).

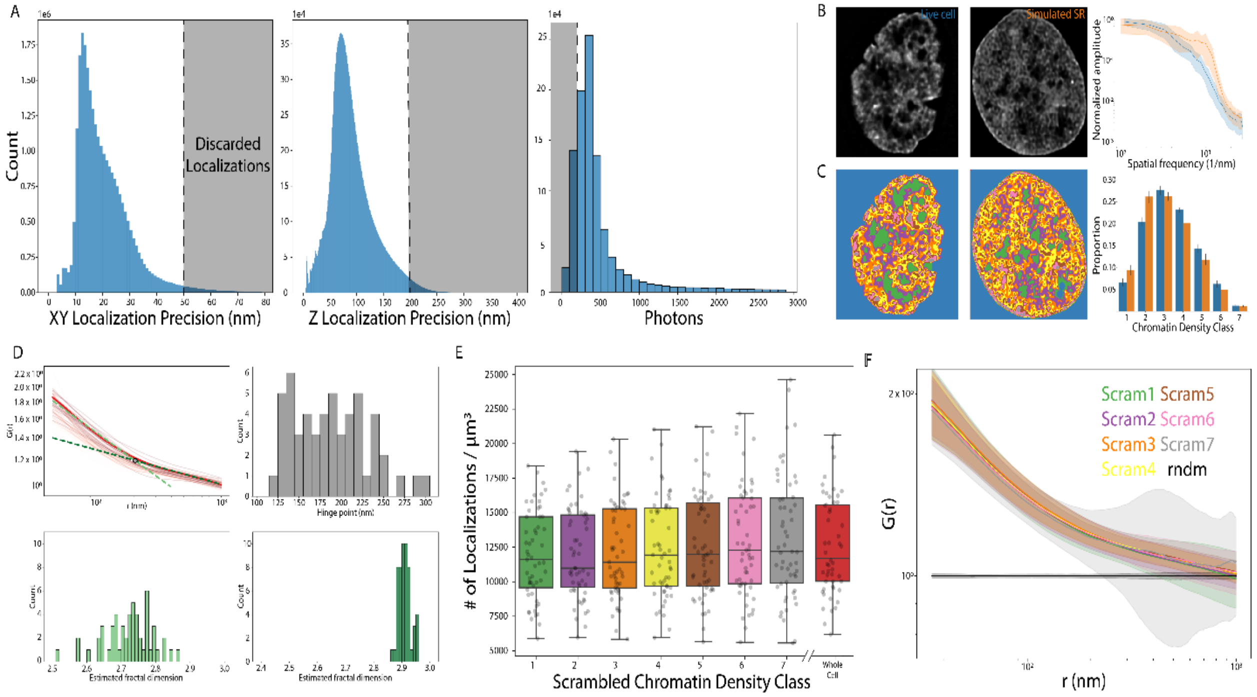


**Figure S4: (A)** The histogram of nucleosomes' lateral (left) and axial (middle) localization error, and the number of photons (right) for fixed-cell HIST single molecule localization microscopy. Shaded areas are discarded by the post-processing localization filter. **(B)** Comparison of image power spectrum between images taken with lattice light sheet microscopy and comparable images generated by convolving the super resolution single molecule localization microscopy dataset with a lattice light sheet microscopy point spread function. Left panel: representative image of deconvolved chromatin structure in live cell taken with lattice light sheet microscopy (same as Figure1E); Middle pane: representative image generated using the single-molecule localizations (same as Figure2D); Right panel: power spectrum of the lattice light sheet microscopy (blue) and convolved single-molecule images (orange). Bar represents mean normalized amplitude and shaded region represents standard deviation. **(C)** The corresponding chromatin density class distribution in (B). **(D)** Estimated fractal dimension based on the pair correlation function (G(r)) calculated over the whole cell. Top left: representative G(r) of whole cells. The thick red indicates G(r) curve for an example cell. The opaque lines represent G(r) curves for other cells. Black circle indicates the hinge point separating the two fitting regimes. The light green line shows the power law fitting over the length scale smaller than the hinge point and the dark green line shows the power law fitting over the length scale larger than the hinge point. Top right: the histogram of hinge points. Bottom left: the histogram of the estimated fractal dimensions smaller than the hinge point. Bottom right: the histogram of the fractal dimensions larger than the hinge point. **(E)** Box plot of nucleosome localization density in different scrambled chromatin classes. The plot convention is the same as figure 1G. **(F)** Pair correlation function of the nucleosome organization for scrambled chromatin classes. The black curve is the G(r) for a random distribution. The plot convention is the same as figure 2K. Data in A are from a single representative cell from 1 biological replicate. Data in B are from n = 20 cells (LLSM) or n = 17 cells (HIST) across a 1 independent biological replicate. Data in D are from n = 54 cells across 3 independent biological replicates. Data in E and F are from n = 54 cells across 3 independent biological replicates.

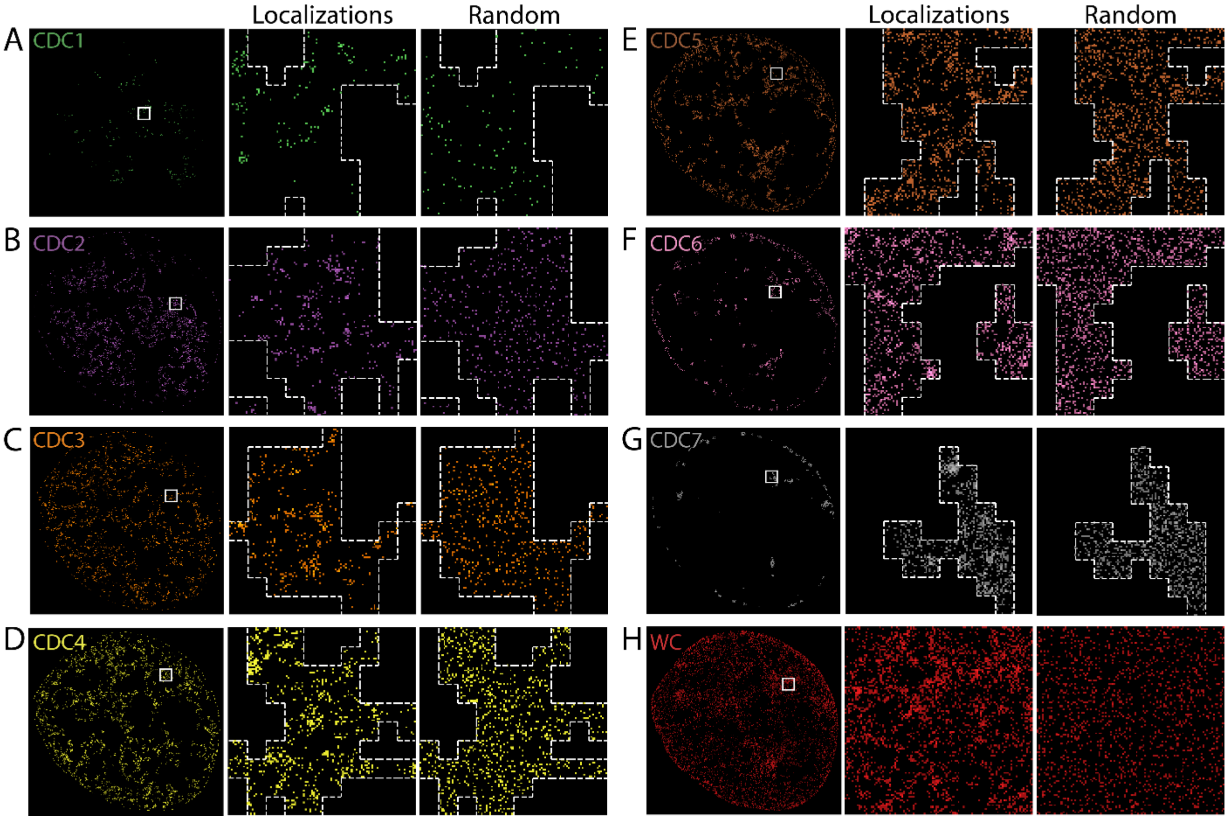


**Figure S5:** Representative comparisons of the experimental and randomized nucleosome distribution in different chromatin density classes. **(A)** Left panel: nucleosome localization associated with chromatin density class 1. Middle panel: zoom-in of the white box in the left panel, the dashed line shows chromatin class 1 boundary. Right panel: random distribution of nucleosome localization with the same density. **(B-G)** Visual comparison between experimental and randomized nucleosome distribution for chromatin density class 2 to 7 respectively, organized similar as (A). **(H)** Visual comparison between experimental and randomized nucleosome distribution of the whole cell, organized similar as (A).

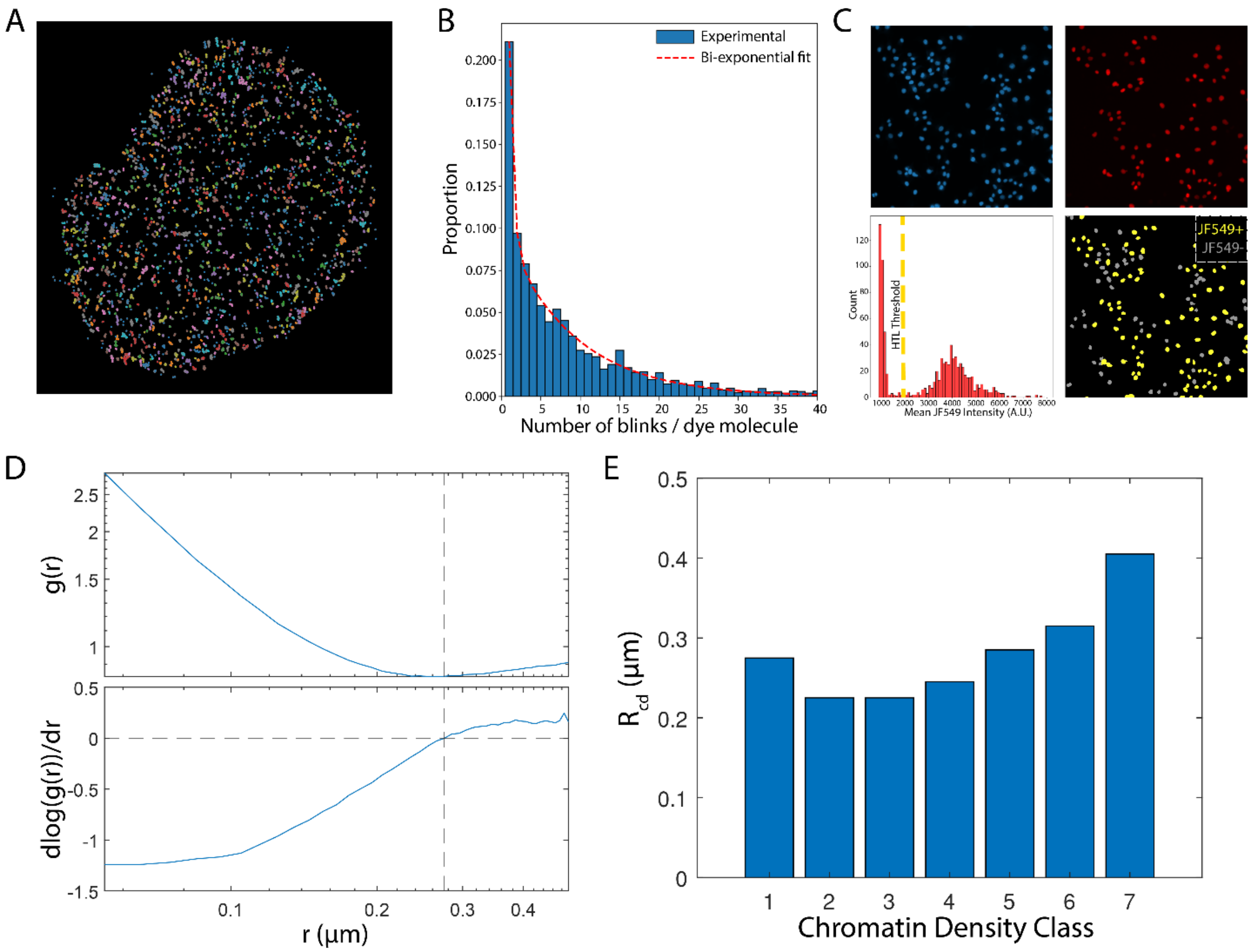


**Figure S6: (A)** Characterization of the blinking properties for JF630B. Nucleosomes are sparsely labelled with the JF630B-HTL dye. All detected localizations are aggregated over 100,000 frames, and the density-based scan method is applied to determine clusters. Each cluster represents the blinking events for a single HaloTag-H2b molecule. **(B)** Histogram of the number of blinks in each JF630b labeled HaloTag-H2b molecule. The dashed line shows a bi-exponential fit to the histogram. **(C)** Epifluorescent images of Cos7-HaloTag-H2b cells labelled with hoechst (blue, top left) and HTL-JF549 (red, top right). The bottom left panel shows the histogram of cell-averaged intensity of JF549, and the yellow dashed line shows the histogram below which cells are considered as not expressing HaloTag-h2b. The bottom right panel shows the segmented nuclei expressing HaloTag-H2b (yellow) and non-expressing (gray) cells. **(D)** Calculation of chromatin domain size used in the biophysical model. The top panel shows the G(r) of a representative cell in log-log scale, and the bottom plot shows the corresponding first derivative. The horizontal dashed line represents when the first derivative reaches zero, and the vertical dashed line is the corresponding radius which is extracted as an estimation of the chromatin domain size. **(E)** The distribution of chromatin domain size for different chromatin density classes. Data in A and B are from n = 13 cells across 2 independent biological replicates. Data in C are from n=846 cells from 1 of 3 biological replicate. Data in E are from n = 54 cells across 3 independent biological replicates.

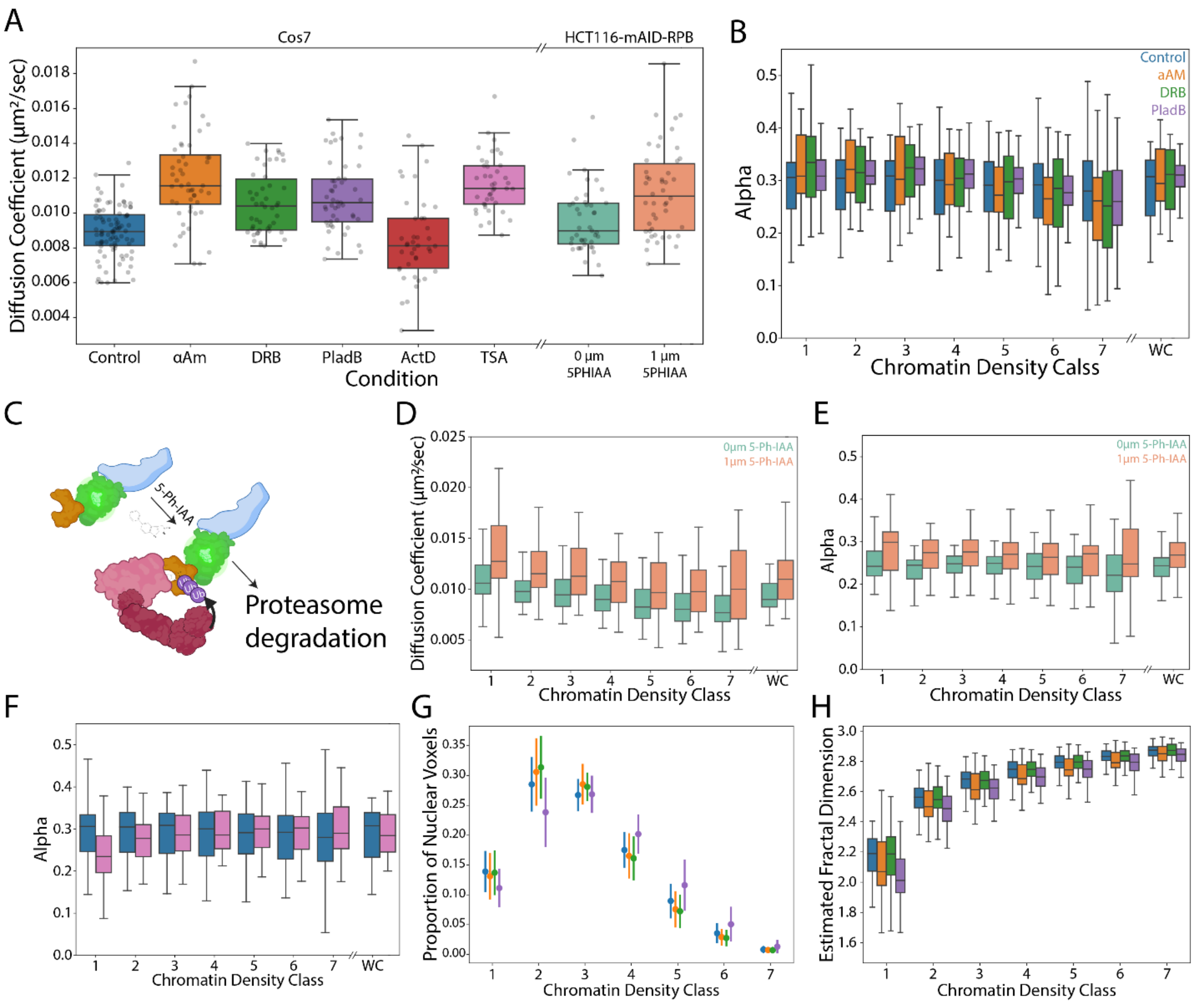


**Figure S7: (A)** Box plot of whole cell averaged diffusion coefficient for different pharmacological perturbations. The plot convention is the same as Figure1G. **(B)** Box plot of MSD anomalous exponent, α, for different chromatin classes under perturbations that inhibit gene transcription. The plot follows the same convention as Figure4B. **(C)** A schematic of conditional knock down of the RNA polymerase II major subunit**. (D)** Box plot of diffusion coefficients of nucleosomes in different chromatin density classes in HCT116-mClover-mAID-RPB+HaloTag-H2b cells under control (green) and after conditional knockdown of RNA Polymerase major subunit with 1 μm 5-Ph-I-AA. **(E)** Box plot of MSD exponent of nucleosomes in different chromatin density classes in HCT116 under control (green) after conditional knock down of RNA polymerase major subunit. **(F)** Box plot of MSD exponent in different chromatin density classes under control (blue) and TSA (pink). **(G)** Distribution of nuclear voxels in different chromatin classes under control (blue), α-amanitin (orange), DRB (green) and PladB (purple). The plot follows the same convention as Figure4G. **(H)** Box plot of the estimated fractal dimension in different chromatin density classes under control (blue), α-amanitin (orange), DRB (green) and PladB (purple). Data in A, B, F, and G are from the same cells and replicates as Figure 4 B-G. Data from D and E are from n = 43 cells (0 μM 5PH-I-AA) and n = 46 cells (1 μm 5PH-I-AA) across 3 independent replicates. Data from H are from n = 54 cells (control), n = 46 cells (aAM), n = 68 cells (DRB), and n = 61 cells PladB across 3 independent replicates.

A

B

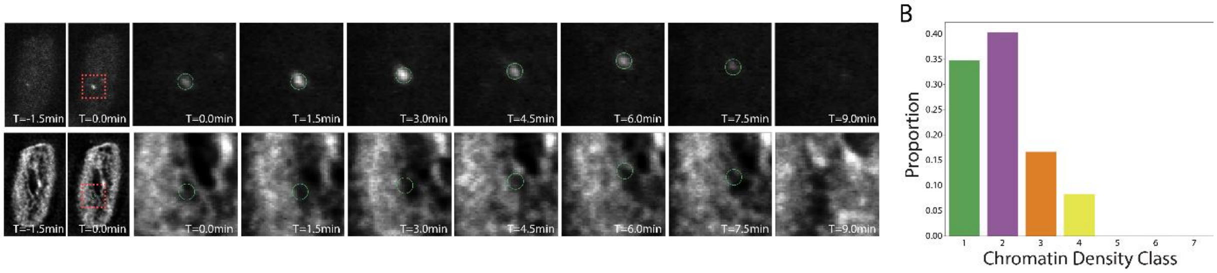


**Figure S8: (A)** A representative example of TFF1 transcriptional burst in MCF7 cells. Top: GFP tagged MS2 coat protein aggregating at TFF1-MS2 transcriptional burst. Bottom: the corresponding chromatin image. **(B)** Histogram of transcriptional burst sites in different chromatin density classes. Data is from n = 105 cells over 3 independent replicates.

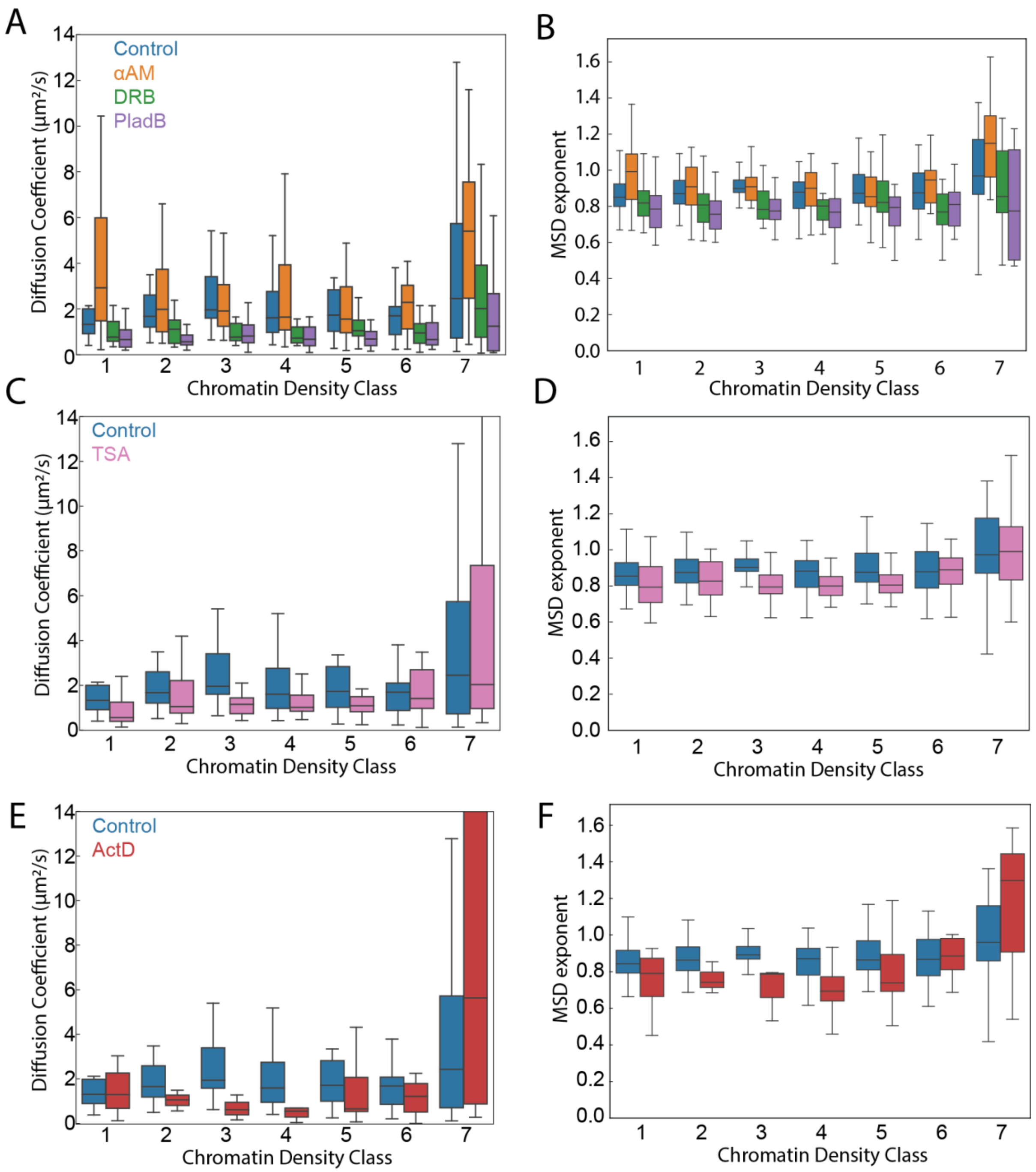


**Figure S9: (A-B)** Box plots of the apparent diffusion coefficient (A) and the anomalous exponent (B) of HaloTag-NLS in different chromatin density classes under control (blue), α-amanitin (orange), DRB (green) and PladB (purple). The plot convention is the same as figure 4B. **(C-D)** Box plots of the apparent diffusion coefficient (C) and anomalous exponent (D) of HaloTag-NLS in different chromatin density classes under control (blue) and TSA (pink). The plot convention is the same as figure 4B. **(E-F)** Box plots of the apparent diffusion coefficient (E) and anomalous exponent (F) of HaloTag-NLS in different chromatin density classes under control (blue) and ActD (pink). The plot convention is the same as figure 4B. Data are from n = 37 cells (control), n = 36 cells (α-amanitin), n = 23 cells (DRB), n = 47 cells (PladB), n = 29 cells (ActD) and n = 42 cells (TSA). Data for control are across four independent replicates, and all other conditions are across three independent replicates.

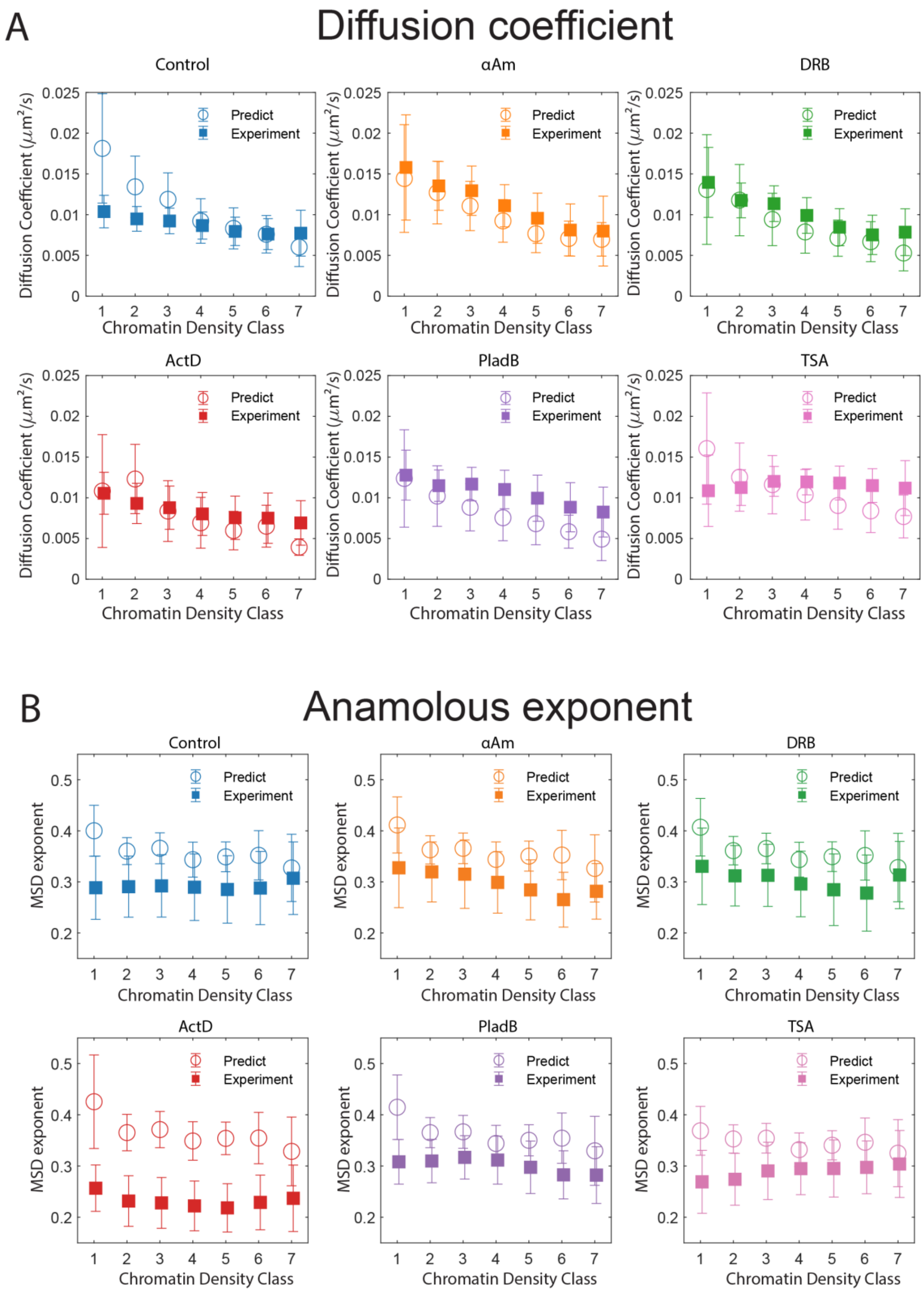


**Figure S10: (A)** Comparison of diffusion coefficients between model (open circles) and experiments (closed squares) under control (blue), α-amanitin (orange), DRB (green), ActD (red), PladB (purple), and TSA treatment (pink). Error bars indicate standard deviations. **(B)** Comparison of the anomalous exponent, figure is organized similarly as (A).
